# Emergence of new function through evolutionary divergence of an intrinsically disordered region

**DOI:** 10.64898/2026.08.25.746950

**Authors:** Carlos A. Elena-Real, Antonia M. Körber, Blaise Gatin-Fraudet, Alexandra Kovinko, Kathrin Motzny, Arbesa Saiti, Adam Lange, Johannes Broichhagen, Sigrid Milles

## Abstract

Intrinsically disordered regions (IDRs) are major drivers of protein functional diversification, yet the molecular features that enable the emergence of new functions within disordered sequences remain poorly understood. FCHO1 and FCHO2 are paralogous pioneer proteins of clathrin-mediated endocytosis that share a conserved domain architecture but perform distinct cellular functions and cannot compensate for each other’s loss. Here we show that functional divergence between these proteins is associated with the acquisition of transient structure within their disordered regions. Using nuclear magnetic resonance spectroscopy, we identify two highly populated α-helical elements in the IDR of FCHO1 that are absent from FCHO2. One of these helices mediates FCHO1 self-association and drives intracellular assembly, whereas FCHO2 lacks this behavior. Introduction of the FCHO1 helix into FCHO2 is sufficient to confer self-association and cellular assembly, demonstrating that a transient structural element embedded within an IDR can act as a transferable functional module. Evolutionary analysis reveals that this helical propensity emerged following duplication of the ancestral FCHO gene and became progressively reinforced during evolution. Despite this divergence, the same region retains a conserved membrane-binding activity in both paralogs. Together, our findings show how acquisition of transient secondary structure within an intrinsically disordered region can generate new molecular behaviors while preserving ancestral functions, providing a mechanism for the functional specialization of paralogous proteins.

## Main Text

Intrinsically disordered regions (IDRs), regions without stable structure, are major drivers of protein functional diversity.^1,2^ Unlike folded domains, IDRs evolve rapidly and can acquire new interaction motifs without disrupting conserved protein architectures.^3^ This evolutionary plasticity is thought to facilitate the emergence of specialized cellular functions and has fueled growing interest in the sequence principles that encode IDR behavior.^4,5^ While considerable progress has been made in understanding how distributed interaction motifs encode collective IDR behaviors such as phase separation,^6–8^ much less is known about how transient secondary structural elements evolve to generate new protein functions.

Gene duplication provides a natural framework for addressing this question. Paralogous proteins frequently retain highly conserved folded domains while diverging within their disordered regions,^9^ suggesting that IDRs may represent privileged substrates for functional specialization. Yet direct molecular examples linking evolutionary changes in IDR structure to the acquisition of distinct cellular functions remain limited.

FCHO1 and FCHO2 constitute an intriguing example of such divergence. These paralogous endocytic proteins, which initiate clathrin-mediated endocytosis,^10^ share the same domain organization and highly conserved folded F-BAR^11^ and μHD^12^ domains (Fig. 1a), yet perform non-redundant biological functions. While FCHO2 is broadly required for constitutive endocytosis,^13^ FCHO1 has specialized roles in immune cells and regulates T-cell receptor internalization.^14^ Notably, the two proteins cannot compensate for each other’s loss,^13,14^ despite their similar domain architectures, suggesting that functional specialization is encoded outside their conserved folded domains.

**Figure 1.**
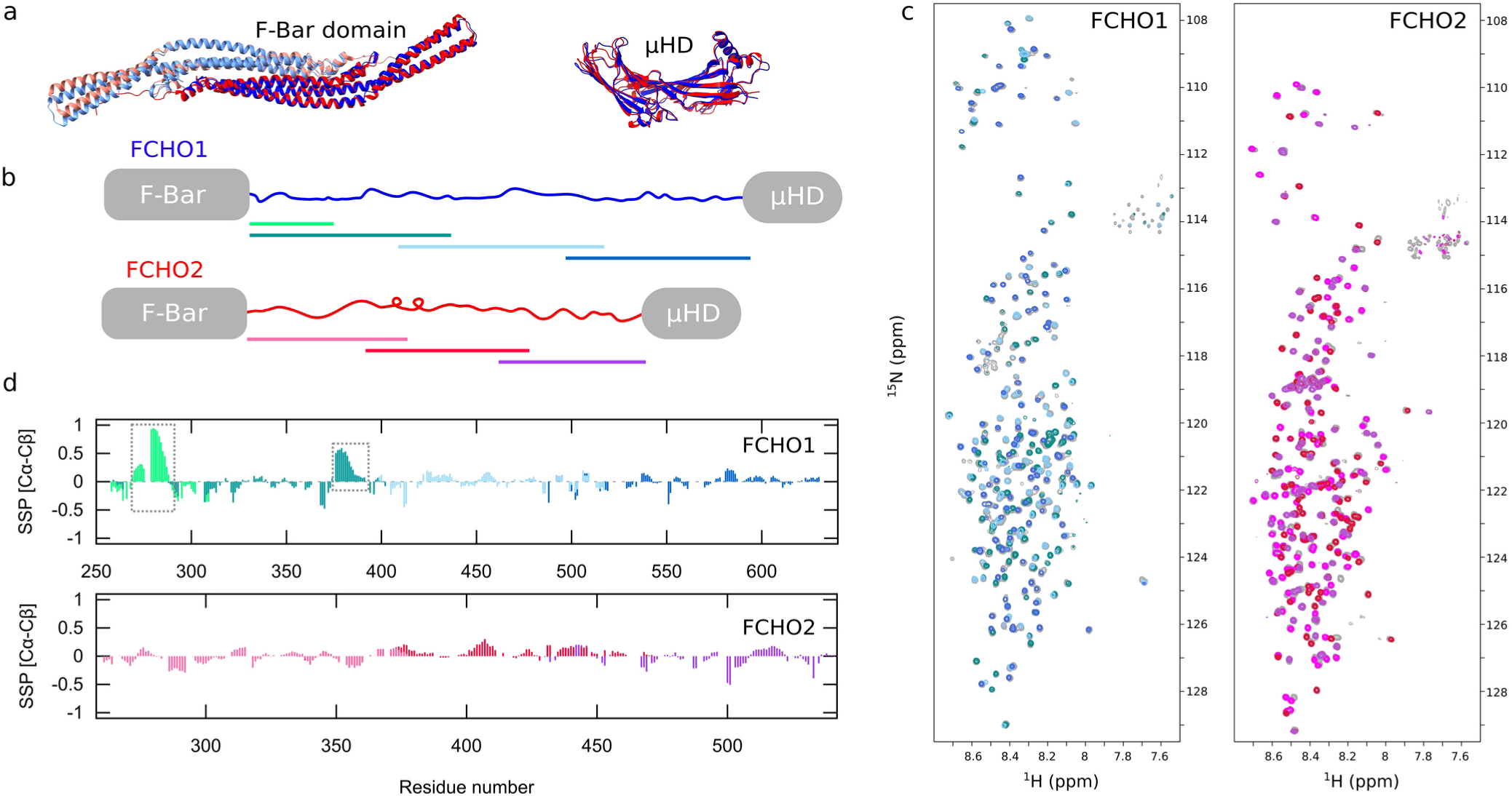
Structural analysis of FCHO1 and FCHO2 IDRs by NMR. **(a)** *Left* – Overlay of ribbon representations of the FCHO2 F-BAR domain dimer crystal structure (red and salmon, PDB: 2V0O^11^) and the AlphaFold2^18^ model of FCHO1 F-BAR domain dimer (blue and light blue). *Right* – Overlay of AlphaFold2 models for the μHD of FCHO1 (blue) and 2 (red). **(b)** Domain architecture of FCHO1 and FCHO2 with the smaller segments used in this study illustrated in different colors. **(c)** ^1^H-^15^N HSQC spectra of FCHO1 and 2 IDRs (grey) and their different segments (colored according to panel b). **(d)** Secondary structural propensities (SSP)^16^ calculated from Cα and Cβ chemical shifts. Positive values correspond to α-helical conformations, negative values to extended conformations, and values near zero to disordered regions. Bars are colored as in panel (b). Dashed boxes indicate helical regions.

Using nuclear magnetic resonance (NMR) spectroscopy, we identify two highly populated α-helical elements within the IDR of FCHO1 that are absent in its paralog FCHO2. We show that one of these helices promotes FCHO1 self-association and drives the formation of intracellular assemblies, whereas FCHO2 lacks both the structural element and the associated cellular behavior. Remarkably, transplantation of this helix into FCHO2 is sufficient to confer self-association and assembly formation, demonstrating that a transient structural element embedded within an IDR can act as a transferable functional module. Evolutionary reconstruction reveals that this helical propensity emerged following duplication of the ancestral FCHO gene and became progressively strengthened during evolution. At the same time, membrane binding mediated by the same region remained conserved between FCHO1 and FCHO2. Together, our findings establish the acquisition of transient structure within an IDR as a mechanism for paralog specialization while preserving ancestral molecular functions.

### α-helical elements sampled by FCHO1’s IDR

Using a divide and conquer strategy,^15^ we produced the full IDR of both FCHO1 and FCHO2, as well as short overlapping segments used for NMR backbone resonance assignment (Fig. 1b). The ^1^H-^15^N heteronuclear single quantum coherence (HSQC) spectra of both proteins displayed the low ^1^H chemical-shift dispersion characteristic of intrinsically disordered proteins (Fig. 1c). We achieved near-complete backbone resonance assignments for both FCHO1 and FCHO2 IDRs (Figs. S1-S7), enabling a detailed analysis of their conformational properties through secondary structure propensity (SSP) calculations based on Cα and Cβ chemical shifts (Fig. 1d)^16^. This analysis, in which fully formed α-helices reach values of 1 and β-sheets values of -1, revealed that FCHO2’s IDR is largely disordered, with SSP values remaining close to zero throughout the sequence (ranging from approximately −0.3 to 0.3). This observation contrasts with previous cryo-electron microscopy structures of the AP2 complex bound to short FCHO2-derived peptides, in which these regions adopt α-helical conformations.^17^

The absence of detectable helical propensity in the isolated protein suggests that FCHO2’s IDR remains predominantly disordered in solution and undergoes coupled folding and binding to the AP2 core complex. In contrast, the SSP profile of the FCHO1 IDR revealed the presence of discrete regions with significant helical propensity. A segment located near the N-terminus of the IDR (residues 270–288) displayed SSP values approaching the theoretical maximum of 1, consistent with the formation of a highly populated and stable α-helix. A second region (residues 376–384) exhibited more moderate SSP values, indicative of a partially populated and dynamic helical conformation. Interestingly, this latter segment overlaps with the predicted AP2-binding site of FCHO1,^17^ suggesting that AP2 recognition may involve distinct structural mechanisms in FCHO1 and FCHO2. Explicit conformational ensembles calculated on the basis of flexible meccano^19^ and ASTEROIDS^20^ (Fig. S8) confirm this observation with population of the first α-helical region to nearly 100%. Together, these results uncover substantial conformational differences between the IDRs of FCHO paralogs. More broadly, they demonstrate that evolutionary divergence within disordered sequences has not merely altered primary sequence composition but has generated distinct structural features that may contribute to the specialized functions of FCHO1 and FCHO2.

### FCHO1 self-associates through α-helical element

To determine whether the sequence-dependent structural differences observed between FCHO1 and FCHO2 IDRs translate into altered backbone dynamics, we performed ^15^N R_1ρ_ relaxation measurements under identical conditions (Fig. 2a). Consistent with its largely disordered nature, FCHO2 IDR exhibits uniformly low and homogeneous relaxation rates across the sequence, indicative of high backbone flexibility and a lack of long-lived structural features. In contrast, FCHO1 IDR displays pronounced site-specific increase in R_1ρ_ values that coincides precisely with regions of high α-helical propensity. The most prominent effect is observed in the N-terminal helix, which exhibits markedly increased relaxation rates, consistent with reduced local mobility due to structural stabilization.^21,22^ The second helical region shows a more moderate increase, reflecting a more transient and dynamically sampled secondary structure. In contrast to FCHO2, FCHO1 showed R_1ρ_ rates with strong concentration dependence, leading to a systematic increase of the rates along the sequence of FCHO1 at 500 μM compared to 200 μM protein concentration. This observation suggests significant self-association of FCHO1, but not FCHO2, which we confirmed using dynamic light scattering (DLS) across a wide concentration range (Fig. 2b). Systematic concentration-dependent increase in particle size, shifting from 8–9 nm of diameter at low concentration (10–50 μM) to 18–20 nm at higher concentrations (600–800 μM), suggests a reversible self-association into higher order assemblies.

**Figure 2.**
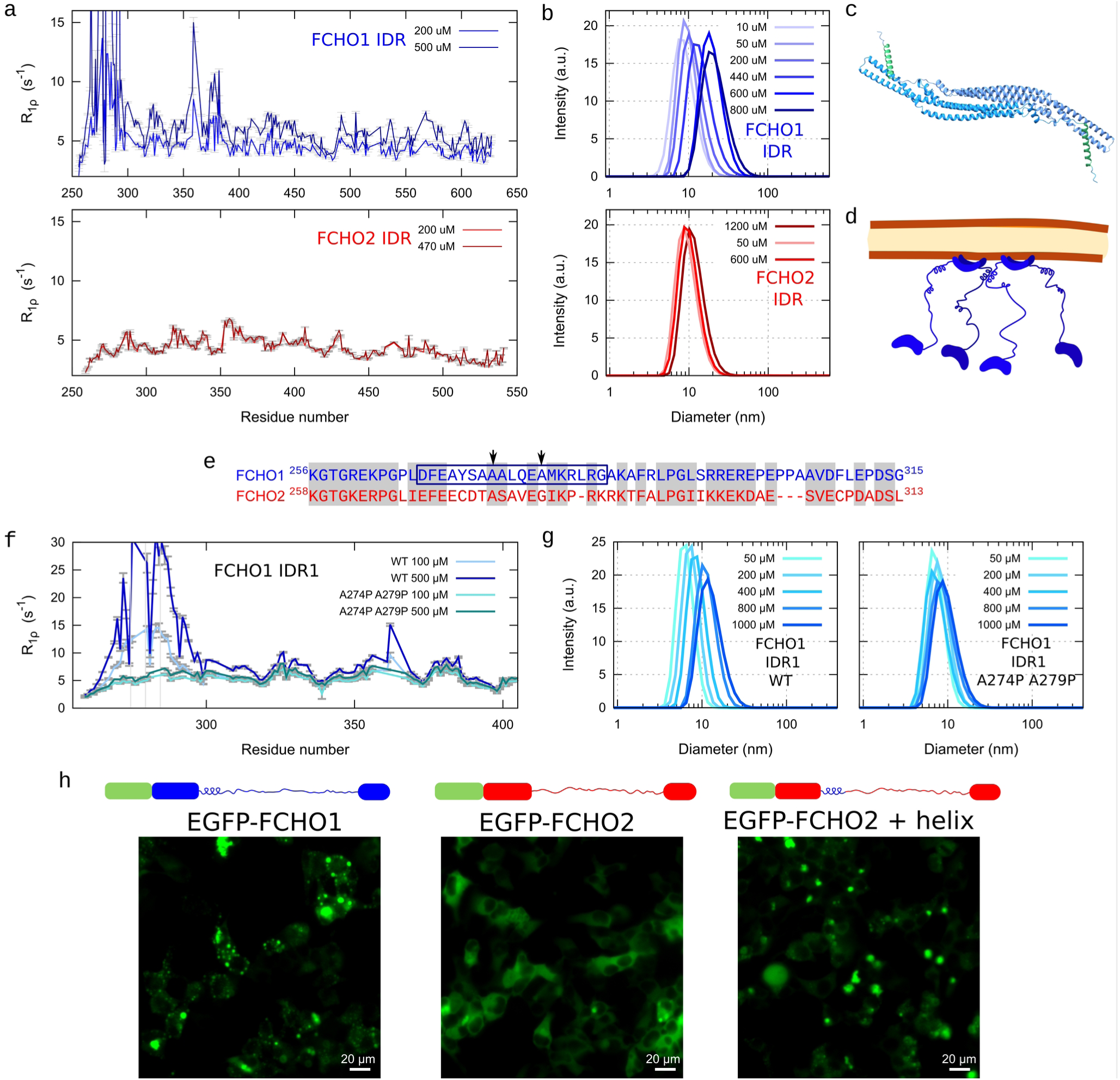
Dynamic analysis of FCHO1 and FCHO2 IDRs. **(a)** ^15^N *R*_1ρ_ spin relaxation measurements of the IDRs of FCHO1 (upper panel) and FCHO2 (lower panel) at two different protein concentrations. **(b)** Dynamic light scattering measurements of FCHO1 IDR (top) and FCHO2 IDR (bottom) upon increasing concentrations. **(c)** AlphaFold2^18,23^ ribbon representation of FCHO1 F-BAR domain dimer (light blue and cyan) including the helix from the IDR (green). **(d)** Schematic model illustrating FCHO1 dimer recruitment through helix–helix interactions. **(e)** Sequence alignment of the N-terminal region of FCHO1 and 2 IDRs. Gray boxes indicate regions of conserved sequence identity or similarity as determined by BLAST^®^.^24^ The black box marks the FCHO1 IDR region forming an α-helix, and black arrows indicate the positions of the Ala→Pro mutations designed to disrupt it. **(f,g)** Dimerization capacity of wt FCHO1-IDR1, and the A274P-A279P mutant, evaluated by ^15^N *R*_1ρ_ spin relaxation (f) and DLS (g) measurements at different concentrations. In the *R*_1ρ_ plots, gray bars indicate the position of A→ P mutations. **(h)** Fluorescence microscopy images of HEK cells transfected with the FCHO variants represented in the top of images. Scale bars are 20 μm.

To identify the region responsible for FCHO1 IDR self-association, we next examined the behavior of the individual IDR segments (Fig. S9). Concentration-dependent dimerization was also observed for the N-terminal fragment (IDR1) (Fig. 2f,g), whereas no such effect was detected for the second and third segments (IDR2 and IDR3) (Fig. S9c). While IDR1 contains the two helical elements identified in the full-length protein, IDR2 includes only the second helical region, which has a smaller helical propensity. The self-association behavior is thus primarily driven by the N-terminal helical region, potentially in cooperation with the adjacent helix.

To disentangle the mechanism underlying FCHO1 IDR assembly, we introduced point mutations designed to selectively disrupt the first α-helix without altering global physicochemical properties such as net charge. The targeted region contains a short alanine-rich segment, including a three-alanine stretch that is absent in FCHO2. We introduced a single alanine-to-proline substitution (A274P) at the center of this helix, and a double mutant (A274P/A279P) to further perturb helical stability (Fig. 2e). We then assessed the impact of these mutations using ^15^N R_1ρ_ relaxation and DLS under identical conditions to the wild-type IDR1 (Fig. 2f,g). The A274P mutation partially disrupted concentration-dependent self-association, as evidenced by reduced increases in hydrodynamic radius and attenuated changes in relaxation rates at high protein concentrations, although a residual concentration dependence remained (Fig. S9d). In contrast, the double mutant (A274P/A279P) strongly suppressed these effects, showing minimal changes in both R_1ρ_ profiles and particle size across the concentration range tested (Fig. 2f,g). Together, these results demonstrate that integrity of the N-terminal helical region is necessary for efficient FCHO1 IDR self-association and that this segment acts as the primary structural determinant of the process. Structural elements within the FCHO1 IDR thus not only modulate local backbone dynamics but also promote concentration-dependent self-association, a property absent in FCHO2. This suggests that sequence-encoded structural motifs within disordered regions can lead to distinct higher-order dynamic behavior of paralogous proteins.

### Transfer of self-association phenotype to FCHO2

Since the N-terminal helix in FCHO1 IDR leading to self-association is situated close to the F-BAR domain, we wondered whether it could participate in the F-BAR dimerization. However, structural models of F-BAR dimers place the IDR helices oriented in opposite directions (Fig. 2c). Therefore, our data support a model in which the FCHO1 IDR introduces an additional, helix-driven interaction mode that cooperates with the F-BAR domain to promote higher-order assembly (Fig. 2d). To address this question, we expressed EGFP-tagged FCHO constructs in HEK293T cells and examined their subcellular distribution by fluorescence microscopy. Consistent with its soluble and dynamic nature, EGFP-FCHO2 exhibited a largely homogeneous cytosolic distribution similar to EGFP alone (Fig. 2h; Fig. S10), in agreement with transient membrane recruitment upon signaling. In contrast, EGFP-FCHO1 formed distinct cytoplasmic accumulations, indicative of higher-order assembly.

While neither deletion of the N-terminal helix nor the F-BAR domain from FCHO1 led to dissolution of the cellular protein assemblies, suggesting that both elements can contribute to assembly stability (Fig. S10), insertion of the FCHO1 N-terminal helical segment (residues 255–288) into the corresponding region of FCHO2, just after the F-BAR domain, was able to reproduce the FCHO1-like intracellular assembly pattern (Fig. 2h). This indicates that this short IDR-encoded element is sufficient to reprogram the assembly phenotype of the FCHO2 paralog.

We therefore propose that the FCHO1 IDR encodes a second layer of self-association that acts in parallel with F-BAR-mediated membrane binding (Fig. 2d). This cooperative mechanism may facilitate intermolecular contacts between FCHO complexes at the membrane, thereby modulating the kinetics and material properties of endocytic site assembly. In contrast to FCHO2, which primarily functions as a membrane recruiter and activator of AP2,^17^ FCHO1 appears to contribute an additional assembly-promoting interface that can tune the organization of the early endocytic network.

### Conserved membrane binding

Several otherwise intrinsically disordered proteins (IDPs) or IDRs have in the past shown to bind to membranes in a helical conformation.^25–29^ Given the close proximity of FCHO1’s N-terminal helix to the membrane, we tested whether it could also be involved in membrane binding.

We prepared lipid vesicles of defined composition, incorporating mixtures of neutral and negatively charged lipids (DOPC:DOPS) to mimic the inner leaflet of the plasma membrane.^30^ The use of large unilamellar vesicles (LUVs), which remain in suspension and are compatible with solution NMR, allowed us to directly probe protein-membrane interactions (Fig. S11). The addition of LUVs induced significant perturbations in the NMR resonances of the FCHO1 IDR. In particular, pronounced line broadening was observed for signals corresponding to the helix spanning residues 270–288, consistent with reduced local mobility upon membrane association and supporting its direct involvement in lipid binding. Indeed, association of FCHO1 with the membranes also led to significant precipitation with the membranes, which we analyzed by solid state NMR spectroscopy. Experiments probing the flexible regions^31^ of FCHO1 demonstrated that, also in the precipitate, the helical region appeared to interact with the membrane, whereas other regions remained flexible (Fig. S12). Fluorescently labeled FCHO1 IDR was efficiently recruited to giant unilamellar vesicles (GUVs) by confocal fluorescence microscopy (Fig. 3a). When we performed the same experiments with FCHO2 IDR, we surprisingly observed a similar phenotype: FCHO2 engaged with membranes through a region with similar distance to the F-BAR domain as FCHO1, with residues in this segment exhibiting a pronounced increase in R_1ρ_ rates consistent with restricted local dynamics due to LUV binding (Figure 3b,c, and Fig. S11). As FCHO1 IDR, fluorescently labeled FCHO2 IDR got efficiently recruited to GUVs (Fig. 3a).

**Figure 3.**
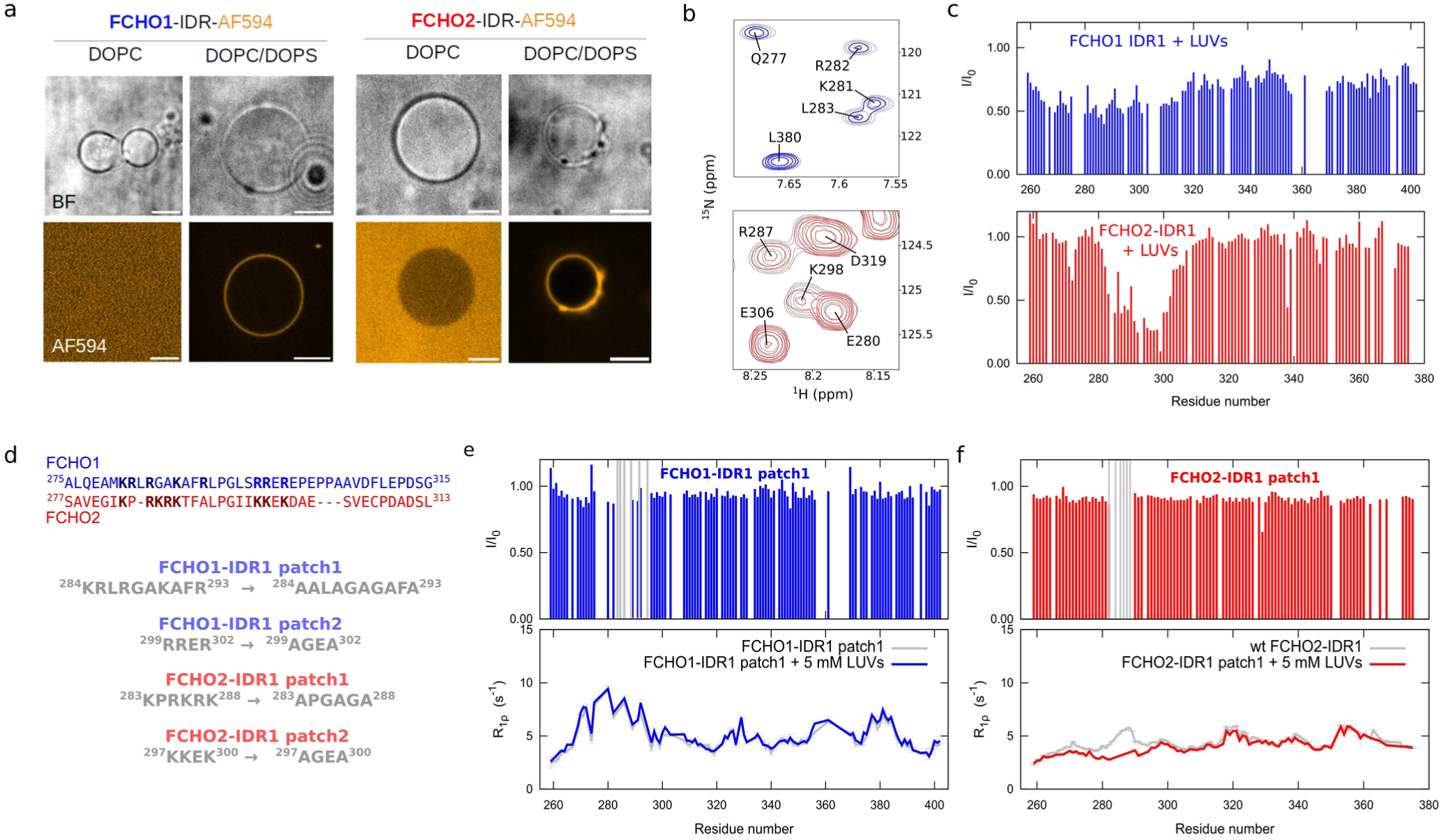
Membrane binding ability of FCHO1 and FCHO2 IDRs. **(a)** Bright field and fluorescent images of GUVs in the presence of fluorescently labeled FCHO1 IDR and FCHO2 IDR. Scale bars are 5 μm. **(b)** Zoom into a ^1^H−^15^N HSQC spectrum of FCHO1 IDR1 (top) and FCHO2 IDR1 (bottom) in the absence (grey) and presence (blue and red) of 120 nm LUVs (DOPC:DOPS 1:1). **(c)** Intensity ratios (I/I_0_) of FCHO1 IDR1 (top) and FCHO2 IDR1 (bottom) in the absence and presence of 120 nm LUVs (DOPC:DOPs 1:1). **(d)** Sequence alignment of FCHO1 and 2 membrane binding regions, highlighting the positively charged residues (top), and the mutants used in this study. **(e,f)** Intensity ratios (I/I_0_) and ^15^N R_1ρ_ spin relaxation of FCHO1 and 2 patch1 mutants in the absence and presence of 115 nm LUVs (DOPC:DOPs 1:1).

To assess whether binding to membranes could induce the formation of a helical element also in FCHO2, we produced micelles from detergent. These 1-palmitoyl-2-hydroxy-sn-glycero-3-[phospho-RAC-(1-glycerol)] (LPPG) micelles interacted with FCHO2 IDR in a similar way as DOPC/DOPS membranes, but they are much smaller than LUVs. This allowed us to maintain NMR signal when FCHO2 was bound to micelles and measure carbon chemical shifts, from which we calculated structural propensities (Fig. S13). The extracted Cα chemical shifts deviated very little from those of the unbound protein, suggesting that FCHO2 binds membranes in its disordered conformation.

Since both FCHO1 and FCHO2 IDRs bind exclusively to negatively charged lipids (Fig. 3a and Fig. S11), we assessed whether binding could be driven through electrostatic interactions. Sequence alignment (by protein BLAST^24^) of the corresponding IDR segment revealed multiple positively charged residues whose positions are partially conserved between FCHO1 and FCHO2, although not always their chemical identity (Fig. 3d). Notably, these residues are organized into two distinct clusters of basic amino acids (hereafter patch 1 and patch 2, containing five and four residues, respectively). The second cluster is particularly striking, consisting of an RRER motif in FCHO1 and a KKEK motif in FCHO2, suggesting that conservation of charge position may be more important than conservation of specific side-chain chemistry. To dissect the contribution of each cluster, we generated four mutants in which charged residues were substituted by alanine or glycine, thereby preserving sequence length while minimizing the introduction of secondary-structure biases (Fig. 3d). We then assessed their ability to interact with negatively charged LUVs using NMR spectroscopy. In FCHO1, mutation of patch 1 completely abolished membrane interaction, whereas mutation of patch 2 produced only a partial reduction in binding (Fig. 3e; Fig. S14). In FCHO2 both patch 1 and patch 2 mutants strongly impaired vesicle association (Fig. 3f; Fig. S14), suggesting a more cooperative contribution of both basic clusters in this paralog, or potentially different membrane affinities of lysine versus arginine in this context.

Interestingly, the pronounced increase in R_1ρ_ rates around the N-terminal helix within FCHO1 IDR was maintained in the protein construct, where patch 1 was mutated and that did not bind to membranes any more, suggesting that helical conformation and membrane binding are also independent in FCHO1. To confirm this, we used the helix-disrupting mutant (A274P/A279P) to evaluate the role of secondary structure in membrane binding. Indeed, disruption of the α-helical conformation did not impair the ability of the FCHO1 mutant to associate with LUVs. NMR intensity ratio profiles were highly similar between wild-type and mutant FCHO1 IDR1, with residues in the 270–288 region remaining strongly affected upon vesicle addition (Fig. S11c).

These results support a model in which FCHO IDR-membrane interactions are primarily governed by electrostatic charge-charge interactions rather than secondary-structure formation. More broadly, they indicate that both FCHO1 and FCHO2 retain a conserved mechanism for binding negatively charged membranes mediated by spatially organized clusters of basic residues. Subtle divergence in the composition and functional contribution of these charge patches between FCHO1 and FCHO2 suggests that evolutionary changes in IDR sequence composition can modulate the quantitative features of membrane recognition while preserving its overall electrostatic basis.

### Evolutionary stabilization of helical traits

Our results indicate that the IDRs of both FCHO paralogs exhibit distinct physicochemical properties, while retaining conserved interactions such as membrane binding within the same protein region. How these functional differences emerged during evolution remains unclear. In non-vertebrate metazoans, a single FCHO gene is present, whereas vertebrates possess two paralogs.^32^ This duplication occurred in the vertebrate lineage, which also saw the emergence of adaptive immunity,^33^ and which may require specialized endocytic regulation, including the internalization of the T-cell receptor in higher vertebrates.

This raises the question of whether the distinctive features of FCHO1 – namely the presence of a stable α-helical element and IDR-mediated self-association – were already present at the time of gene duplication or emerged later during vertebrate evolution. To address this, we performed a multiple sequence alignment of the FCHO1 region corresponding to the high helical propensity segment (residues 270–288) across representative vertebrate species (Fig. 4a). Remarkably, this α-helical sequence is nearly invariant among mammalian species, whereas it is less well conserved in zebrafish and frog, with chicken displaying an intermediate degree of conservation. Importantly, the positively charged residues implicated in membrane binding are preserved across all analyzed species. Although direct experimental assessment of oligomerization across species remains challenging, structural predictions can provide insight into the propensity of these regions to form helices. Given that AlphaFold2^18^ reliably predicts the helical structure of the human FCHO1 270–288 segment, we extended this analysis to orthologous sequences from different vertebrates (Fig. 4b). Consistent with the sequence conservation pattern, mammalian orthologs are predicted to form α-helices of similar length and stability as the human protein. In contrast, non-mammalian vertebrates display reduced but still detectable helical propensity, suggesting that this region may have originated as a weak, dynamic structural element that progressively stabilized during vertebrate evolution. This supports a model in which a weak helical tendency emerged early in FCHO1 evolution and was subsequently refined, potentially under selective pressure to enhance the self-association and membrane-proximal assembly properties described here. Notably, the emergence of a stable α-helical element adjacent to an F-BAR domain appears to be specific to FCHO1, as such architecture is not observed in other F-BAR containing proteins (Fig. 4c).

**Figure 4.**
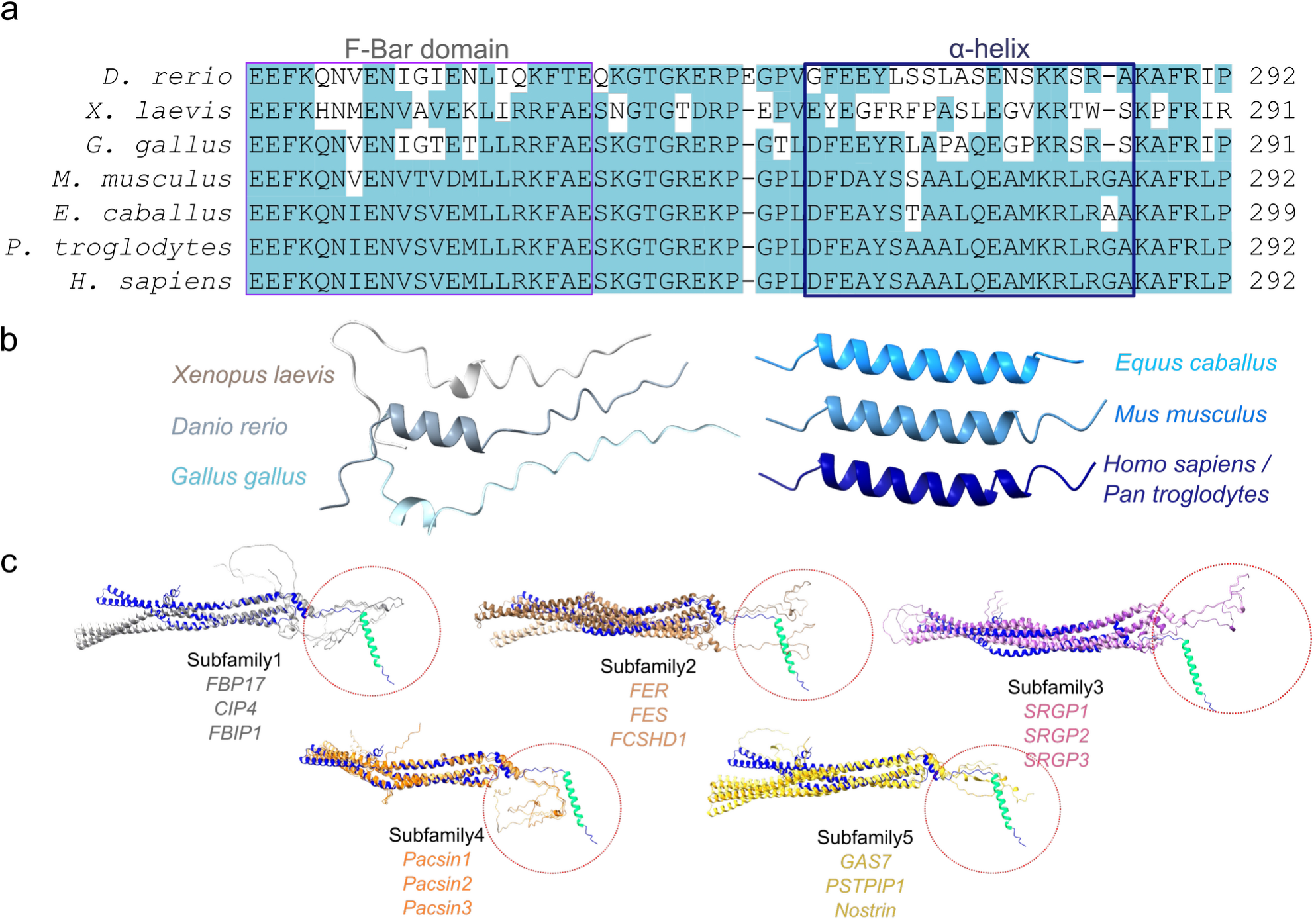
Evolution of FCHO IDR N-terminus. **(a)** Multiple sequence alignment (Clustal Omega^34^) of FCHO1 IDR N-terminal stretch from 7 different vertebrates. Boxes indicate the F-BAR domain and the location of the α-helix discovered in this study. Blue color indicates sequence similarity with the human sequence. **(b)** AlphaFold2 models of the IDR stretches corresponding to the human helix. **(c)** AlphaFold models of extended F-BAR domains of 15 members of different families of F-BAR domain-containing proteins, overlapped with the model of the extended F-BAR domain of FCHO1. F-BAR domain of FCHO1 is in blue and its IDR populating a helical conformation in green. Other F-BAR domains and their tails of similar lengths are indicated in different colours.

Together, these observations support a model in which FCHO1 and FCHO2 diverged through step-wise remodeling of their IDRs, leading to the emergence of distinct, lineage-dependent interaction capabilities that were not present in the ancestral FCHO protein.

## Conclusions

Our findings provide a framework for understanding how IDRs evolve new molecular functions. Rather than replacing ancestral activities, the same short segment of the FCHO disordered region has accumulated distinct functional features during evolution: conserved positively charged residues maintain membrane binding, whereas subtle sequence changes generated new interaction modules, including a transient α-helix that promotes self-association – an interaction mode that is quite different to previous transient interaction sites identified in endocytic IDRs.^15,35,36^ The emergence of this function required only limited modifications to local sequence composition, illustrating how IDRs can acquire new molecular behaviors without the structural constraints imposed by folded domains.

This evolutionary plasticity offers an explanation for why FCHO1 and FCHO2, despite their similar domain organization, are not functionally interchangeable^13,14^. More generally, it suggests that paralog specialization can arise through the progressive functionalization of disordered regions, enabling increasingly specialized cellular pathways while preserving core ancestral activities. Beyond its evolutionary implications, our work highlights transient structural elements as modular and engineerable features of IDPs, expanding current paradigms for understanding and designing IDR function^6–8^ beyond sequence composition alone.

## Methods

### Cloning

pET28a-FCHO-6His vectors for bacterial recombinant expression were purchased from Twist Bioscience, using *Nco*I and *Xho*I as restriction enzymes. FCHO2-IDR3 was cloned with an N-terminal GB1 solubility tag into a pET28-6His-GB1 vector using *Nde*I and *Not*I restriction enzymes.^15^ The constructs FCHO1-IDR_255-624_, FCHO1-IDR1_255-405_, FCHO1-IDR1short_255-306_, FCHO1-IDR2_363-514_, FCHO1-IDR3_475-624_ were designed using FCHO1 UniProt ID O14526, while for the constructs FCHO2-IDR_258-542_, FCHO2-IDR1_258-373_, FCHO2-IDR2_333-468_ and FCHO2-IDR3_428-542_ the FCHO2 UniProt ID Q0JRZ9 was used. FCHO1-IDR1 A274P and A274P/A279P mutants were constructed using site-directed mutagenesis.

For the fluorescent expression in human cells, full length FCHO proteins and their chimeras were purchased from Twist Bioscience within pTWIST vectors, and subsequently cloned into pEGFP-C3 using *Bgl*II and *Sal*I restriction enzymes.

The primer list for all cloning performed can be found below:

### Restriction enzyme cloning

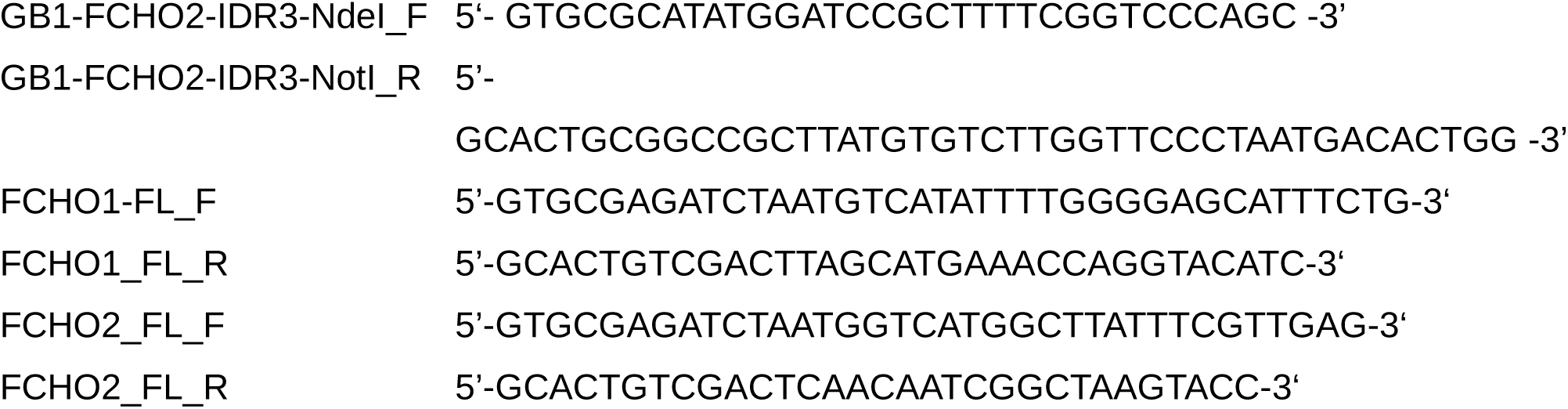

### Site directed mutagenesis

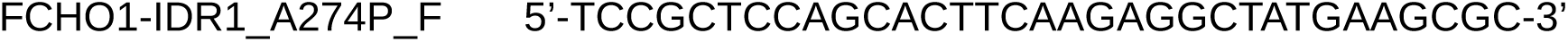
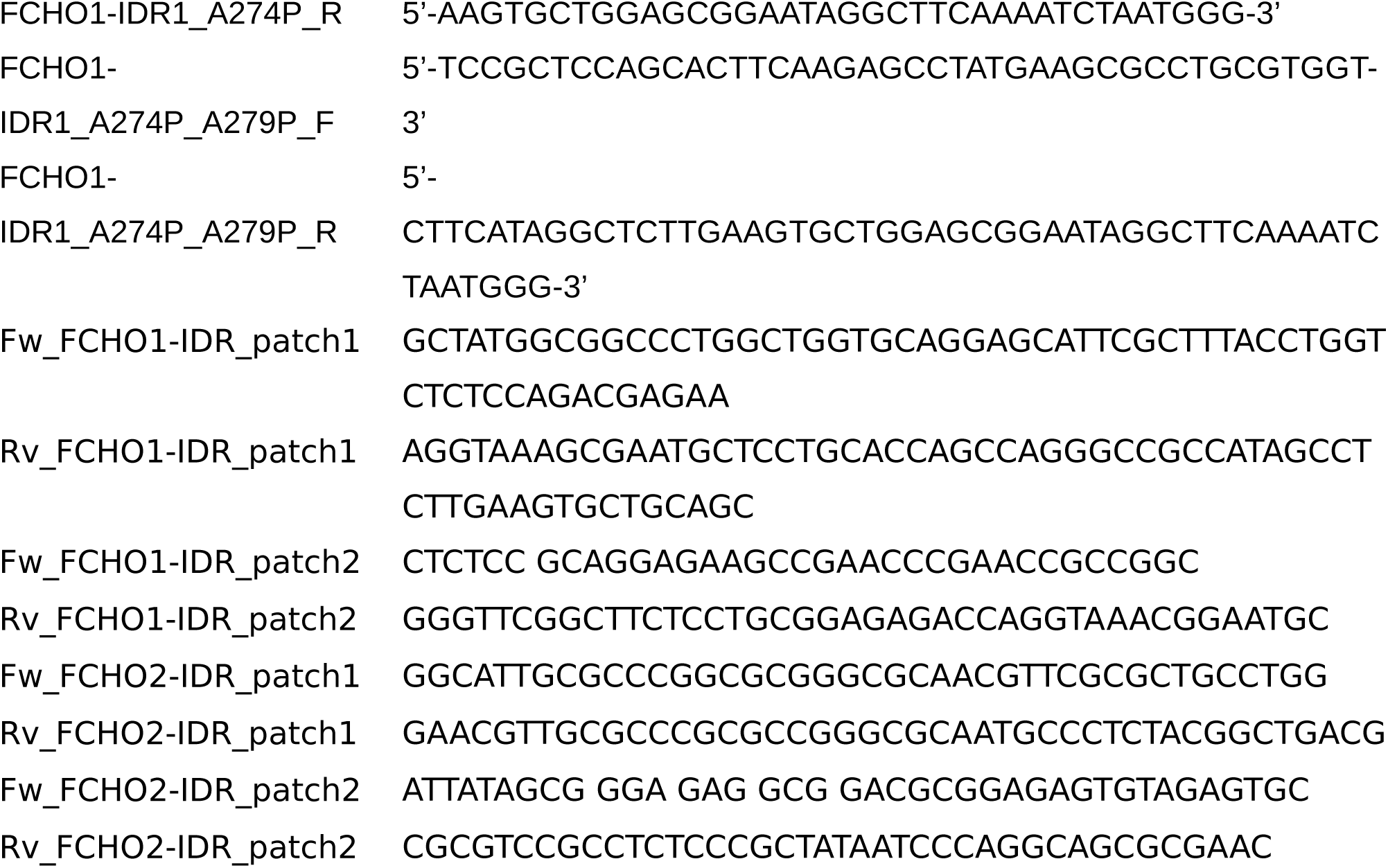

### Protein expression and purification

Recombinant expression of proteins was performed using *E. coli* Rosetta (DE3) strain and grown in LB medium with 30 mg/L Kanamycin and 30 mg/L Chloramphenicol at 37 °C. Expression was induced with 1 mM isopropyl-β-D-thiogalactopyranoside (IPTG) when the optical density at 600 nm reached ∼0.6, then cultures were incubated at 20 °C overnight. For isotope labeling (^15^N, ^13^C), M9 minimal medium supplemented with 1 g/L ^15^NH_4_Cl and/or with 2 g/L ^13^C-glucose was used.

Cells were lysed by sonication at 4 °C in 20 mM Tris (pH 8.0), 150 mM NaCl, with Roche Ethylenediaminetetraacetic Acid (EDTA)-free protease inhibitor cocktail (Sigma-Aldrich Chemie GmbH). For the full IDR constructs of both FCHO1 and 2, 4 M urea was additionally contained in the buffer. All proteins were purified by affinity chromatography using a hand-packed nickel column and 400 mM imidazole to ensure protein elution. Then, size-exclusion chromatography was performed using a Superdex 75 column equilibrated in NMR buffer (50 mM Sodium phosphate pH 6.0, 150 mM NaCl, and 2 mM dithiothreitol (DTT)). Finally, proteins were concentrated and concentration was determined by absorbance at 280 nm using extinction coefficients determined by Expasy Protparam^37^.

### Solution NMR spectroscopy

All NMR experiments were performed at the Leibniz-Forschungsinstitut für Molekulare Pharmakologie (FMP), Berlin, Germany. Two spectrometers, equipped either with room-temperature-(750 MHz) or cryo-probes (600 MHz), were used in this study for solution NMR. All the solution NMR experiments were performed in NMR buffer (50 mM Sodium phosphate pH 6.0, 150 mM NaCl, and 2 mM DTT) with 10% D_2_O, at 10 °C. Spectra were acquired with TopSpin 3.5 and 4.4.1 (Bruker), processed with NMRPipe^38^, using qMDD^39^ for non-uniformly sampled assignment spectra, and analyzed with CCPN^40^. BEST-TROSY-HSQC spectra were used to determine amide (^1^HN and ^15^N) chemical shifts, while BEST-TROSY triple resonance experiments were acquired to correlate CO, Cα, Cβ resonances and achieve assignments.^41^ Specifically, HNCO, HNCOCA, HNCA, iHNCA, HNCOCACB, iHNCACB, HNCANNH and HNCOCANNH experiments were used to assign the short, overlapping constructs. Experiments included in NMRlib^42^ were used. Cα and Cβ chemical shifts were used to calculate secondary structure propensities^16^, which were compared with random coil values from refDB.^43^

^15^N *R*_1ρ_ spin relaxation rates^44^ were acquired at 600 MHz ^1^H Larmor frequency with a spin-lock field set to 1500 Hz, and using 6 delays, between 10 and 230 ms, to sample the decay of magnetization. The peak intensity relative to the delay time was fit with an exponential curve to determine the relaxation rates, with errors derived from the experimental uncertainty.

### Conformational ensemble of FCHO1 IDR1

A conformational ensemble was calculated from FCHO1 IDR1 using a combination of the statistical coil generator flexible meccano^19^, and the genetic algorithm ASTEROIDS^20^. From a statistical coil ensemble of 10,000 conformers, 200 conformations were selected, which together best described the HN, N, CO, Cα and Cβ chemical shifts of FCHO1 IDR1. A new ensemble of 8500 conformers was generated using the Φ and ψ angles from the previous selection, to which 1500 conformations from the previous pool of conformers were added. Another selection of 200 conformations was performed with the new pool. The process was iterated 4 times. Ensemble-averaged chemical shifts were generated using SPARTA^45^, and secondary chemical shifts were calculated based on RefDB^43^.

### Solid-state NMR spectroscopy

Uniformly ^15^N-labeled FCHO1 IDR1 (300 μM) was mixed with 4 mg LUVs at pH 5, inducing vesicles precipitation. The sample was packed into a 1.3 mm zirconia MAS rotor. Prior to closing the rotor, 0.2 μL of a saturated solution of DSS (4,4-dimethyl-4-silapentane-1-sulfonic acid sodium salt) in D₂O was added.

All solid-state NMR experiments were performed on a Bruker 600 MHz spectrometer equipped with a 1.3 mm triple-resonance (HCN) MAS probe. The magic-angle spinning frequency was set to 55 kHz, and the sample temperature was maintained at 293 K.

A two-dimensional hNH INEPT correlation spectrum was acquired with spectral widths of 40 ppm in the ^1^H dimension and 23 ppm in the ^15^N dimension. The INEPT transfer delays were set to τ₁ = 2.5 ms and τ₂ = 2.7 ms, respectively, and were optimized to maximize the signal intensity. Hard 90° pulses were calibrated on each channel, corresponding to radiofrequency powers of 5.8 W (100 kHz) for ^1^H and 12.7 W (35.7 kHz) for ^15^N. A recycle delay of 1.5 s was used, with 384 scans per t₁ increment. The time-domain data consisted of 2048 complex points in the ^1^H dimension and 96 complex points in the ^15^N dimension.

The spectra were processed using Bruker TopSpin and chemical shifts were referenced internally to DSS. Data analysis was performed using CCPN^40^.

### Dynamic light scattering

Dynamic light scattering experiments were performed to evaluate the dimerization state of IDR constructs from FCHO1 and 2, as well as the A274P and A274P/A279P mutants. Measurements were carried out at different protein concentrations ranging from 50 to 1000 μM in NMR buffer at 25 °C. A Zetasizer Nano ZS instrument was used (Malvern Instruments).

### Fabrication of membrane vesicles

Large and giant unilamellar vesicles (LUVs and GUVs, respectively) were prepared using 1,2-dioleoyl-sn-glycero-3-phosphocholine (DOPC) and 1,2-dioleoyl-sn-glycero-3-phospho-L-serine (DOPS) (Avanti Polar Lipids), supplied in chloroform.

For LUV preparation, the lipids were mixed (DOPC and DOPS in a 1:1 molar ratio, final 5 mM lipids concentration) in a round-bottom glass vial followed by evaporation of the solvent using a rotary evaporator. After hydration of the dried lipid film with NMR buffer (pH 5.0 or 7.0), the lipid vesicle solution was extruded first through 400 nm, then through 100 nm filters, each 35 times, to obtain uniform lipid vesicles with a diameter of around 100 nm. Approximate diameters determined by DLS of the different vesicle preparations are indicated in the figures. Finally, the lipid vesicle solution was stored in glass vials at 4 °C.

GUVs were prepared by electroformation using 100 nmol of lipids per sample. The lipids were mixed in chlororform (DOPC and DOPS in a 1:1 molar ratio) and evenly applied as thin layers onto preheated titanium plates (50 °C). After the plates were assembled with parafilm as insulating layers, they were dried under high vacuum for 1 h to remove traces of solvent. The electroformation chambers were sealed with parafilm and filled with sucrose buffer (250 mM sucrose, 1 mM NaN_3_, 280 mOsmol/kg). An alternating electric field of 10 Hz, rising from 0.02 V to 1.1 V, was applied for 3 h at 55 °C. The GUVs were stored at room temperature until they were used for microscopy assays.

### Fluorescence labelling

FCHO1 IDR and FCHO2 IDR were labeled with AZDye 594 (AF594) Maleimide (Vector Laboratories, Inc., CA, USA) essentially as described previously.^46^ Briefly, protein samples were incubated in labeling buffer (50 mM Na-phosphate pH 7.0) supplemented with 10 mM Dithiothreitol (DTT) over night at 4°C. The reducing agent was then washed out using extensive dialysis into labeling buffer. Dry AZDye 594 Maleimide was then dissolved in water-free dimethyl sulfoxide (DMSO) and added to the protein at a molar excess of 5 times under gentle agitation. The labeling reaction was allowed to proceed for one hour at room temperature and continued at 4°C over night. Labeled protein was then separated from free dye using size exclusion chromatography on a SEC70 column (BioRad). Protein purity was assessed by SDS-PAGE and pure fractions were concentrated. Protein concentrations were determined based on the absorbance by the fluorophore.

### Fluorescence microscopy

GUVs were transferred to Poly-L-Lysine coated 8-well imaging slides (ibidi) and diluted 1:3 with imaging buffer (250 mM glucose, 5.8 mM KH_2_PO_4_, 5.8 mM K_2_HPO_4_, 300 mOsmol/kg). After GUVs settled at the bottom of the imaging slide, fluorescently labeled proteins were added (final concentration 1 μM). Confocal imaging was performed with a Nikon CSU-X1 spinning disc confocal microscope equipped with a 60x oil immersion objective. The setup was controlled by the imaging software NIS (Nikon). Microscopy images were processed using the software ImageJ/Fiji^47^.

### In-cell experiments

HEK293T cells were cultured in growth medium (DMEM, Glutamax, 4.5 g Glucose, 10% FCS, 1% Penicillin-Streptomycin) at 37 °C and 5% CO_2_. 50,000 cells per well were seeded on 8-well µL slides (ibidi) previously coated with poly-L-lysine. The next day, 400 ng DNA was transfected using 0.8 µL Jet Prime reagent in 40 µL Jet Prime buffer (VWR) per well/plasmid. Medium was exchanged against antibiotic-free media before the transfection mix was pipetted on the cells. After 4 hours incubation at 37 °C and 5% CO_2_, medium was exchanged against growth media. After additional 24 hours, cells were imaged after one washing step in Fluorobrite (Thermo Fisher) at 37 °C and 5% CO_2_ using a Ti-E Nikon epifluorescence microscope equipped with pE4000 (cool LED), Penta Cube (AHF 66-615), 60x oil NA 1.49 (Apo TIRF Nikon) and imaged on SCMOS camera (Prime 95B, Photometrics) operated by NIS Elements (Nikon). For excitation the following wavelength and emission the following emission filter were used: λ = 470 nm with 525/40 (Semrock). Images were processed using FIJI.

### Evolutionary analysis

Phylogenetic studies were performed by multiple sequence alignment using Clustal O (1.2.4)^48^. For that, the FCHO1 sequences were obtained from UniProt for *H. sapiens* (O14526), *P. troglodytes* (H2QFR7), *M. musculus* (Q8K285), *E. caballus* (A0A3Q2HZ70), *G. gallus* (A0A8V1AH28), *X. laevis* (A0A1L8HXQ9) and *D. rerio* (E7FBF7).

For the search of a helix connected to F-BAR domain, AlphaFold2 models for three members of the five F-BAR domain proteins subfamilies^49^ were obtained from UniProt. FBP17 (Q96RU3), CIP4 (Q15642), FBIP1 (Q5T0N5), FER (P16591), FES (P07332), FCHSD1 (Q86WN1), SRGP1 (Q7Z6B7), SRGP2 (O75044), SRGP3 (O43295), Pacsin1 (Q9BY11), Pacsin2 (Q9UNF0), Pacsin3 (Q9UKS6), Gas-7 (O60861), PSTPIP1 (O43586) and Nostrin (Q8IVI9) models were aligned in Chimera X^50^ with the AlphaFold2 model of the F-BAR domain of FCHO1 including IDR1short (O14526).

## Data availability

Backbone resonance assignments have been deposited in the Biological Magnetic Resonance Data Bank with the following accession codes: 53887, 53888, 53889, 53890, 53891, 53892 and 53893.

## Acknowledgements

We thank all members of the Milles group for fruitful discussions and critical proofreading. We thank M.R. Jensen for providing the pET28-6His-GB1 plasmid, M. Blackledge and M.R. Jensen for providing NMR analysis scripts, P. Müller for the GUV machine, M. Bashawat for an introduction to fabricating GUVs, P. Schmieder, N. Trieloff and M. Beerbaum for technical assistance on the NMR spectrometers, and T. Soykan, M. Bieck and M. Lehmann from the Imaging Facility of FMP for technical support.

## Author contributions

S.M. and C.A.E.R. conceived the study. C.A.E.R., A.M.K., K.M. and A.S. made samples. C.A.E.R. and A.M.K. recorded solution NMR experiments. A.K. and A.L. recorded and interpreted ssNMR data. C.A.E.R. and A.M.K. performed membrane binding assays and imaging of GUVs. C.A.E.R. performed DLS data. B.G.F. and J.B. performed and analyzed in-cell data. C.A.E.R, A.M.K. and S.M. analyzed and interpreted all data. C.A.E.R. and S.M. wrote the paper with input from all authors.

## Materials & Correspondence

Correspondence and material requests should be addressed to Sigrid Milles

