## Supporting Information for "Emergence of new function through evolutionary divergence of an intrinsically disordered region"

[1]Leibniz-Forschungsinstitut für Molekulare Pharmakologie, Robert-Rössle-Straße 10, 13125 Berlin, Germany.

*****

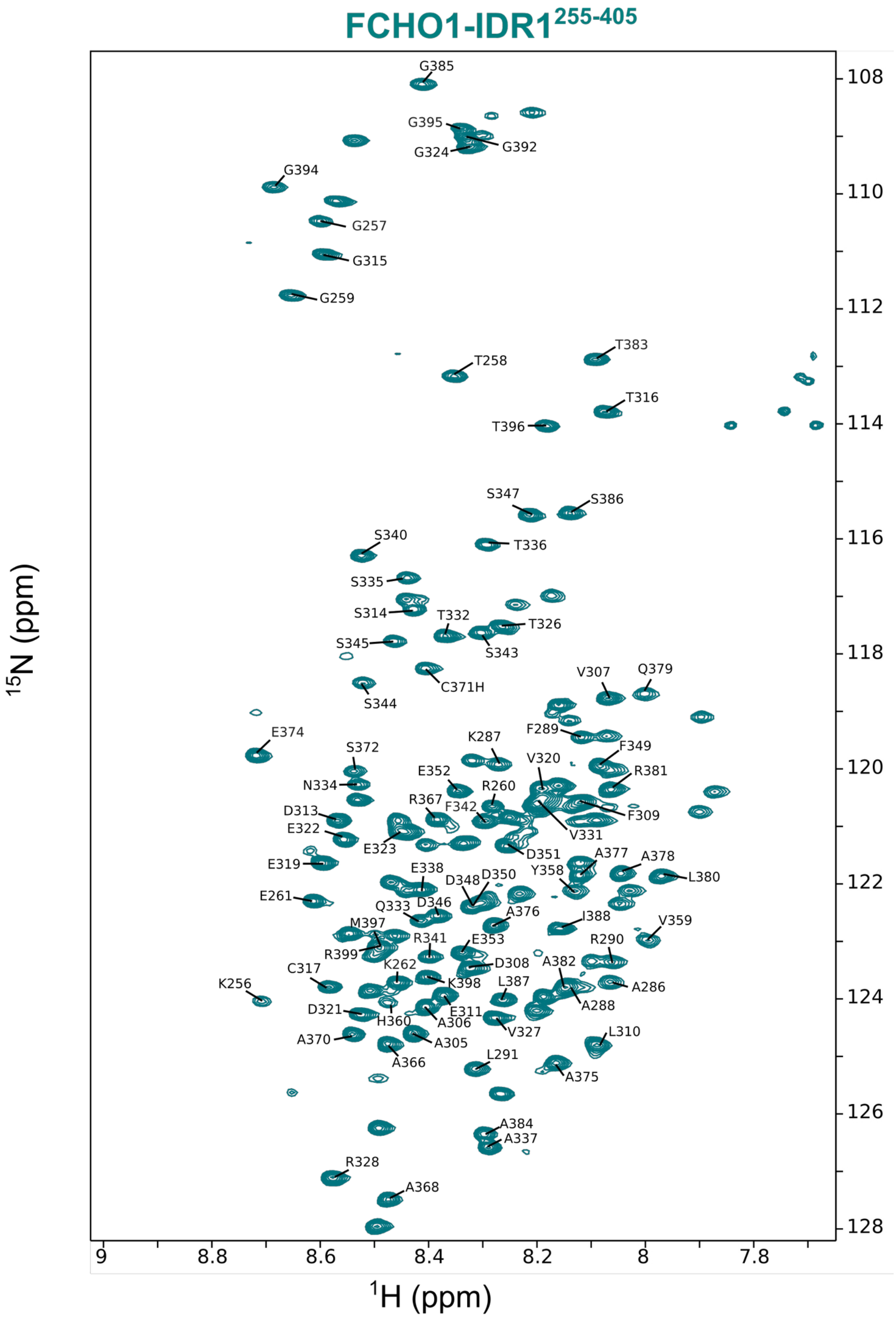


**Fig. S1. FCHO1-IDR1 assignment.** The $^{1}H$-$^{15}N$ HSQC spectra of FCHO1-IDR1 construct showing the backbone resonance assignments as one letter amino acid code.

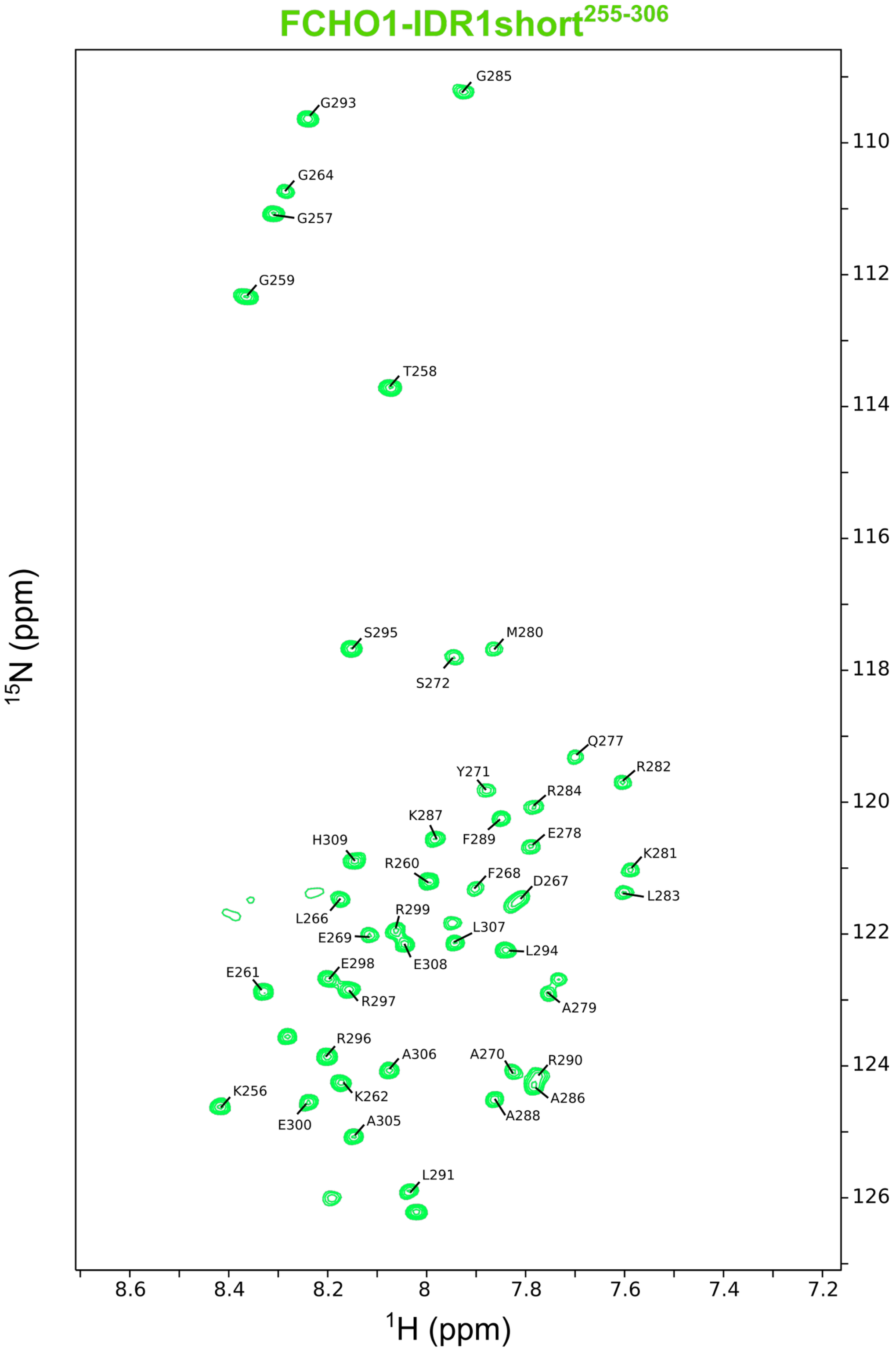


**Fig. S2. FCHO1-IDR1short assignment.** The $^{1}$H-$^{15}$N HSQC spectra of FCHO1-IDR1short construct showing the backbone resonance assignments as one letter amino acid code.

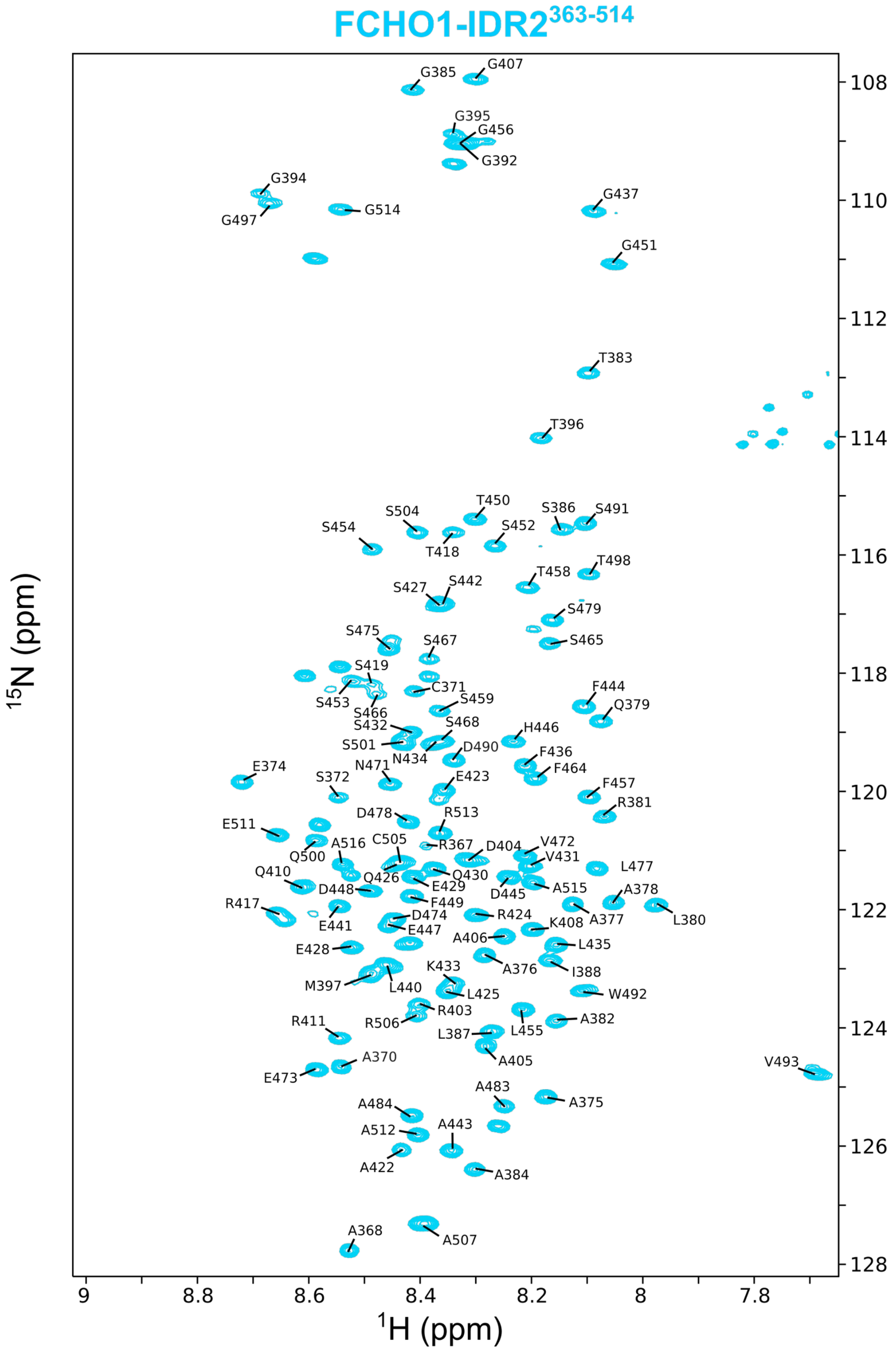


**Fig. S3. FCHO1-IDR2 assignment.** The $^{1}H$-$^{15}N$ HSQC spectra of FCHO1-IDR2 construct showing the backbone resonance assignments as one letter amino acid code.

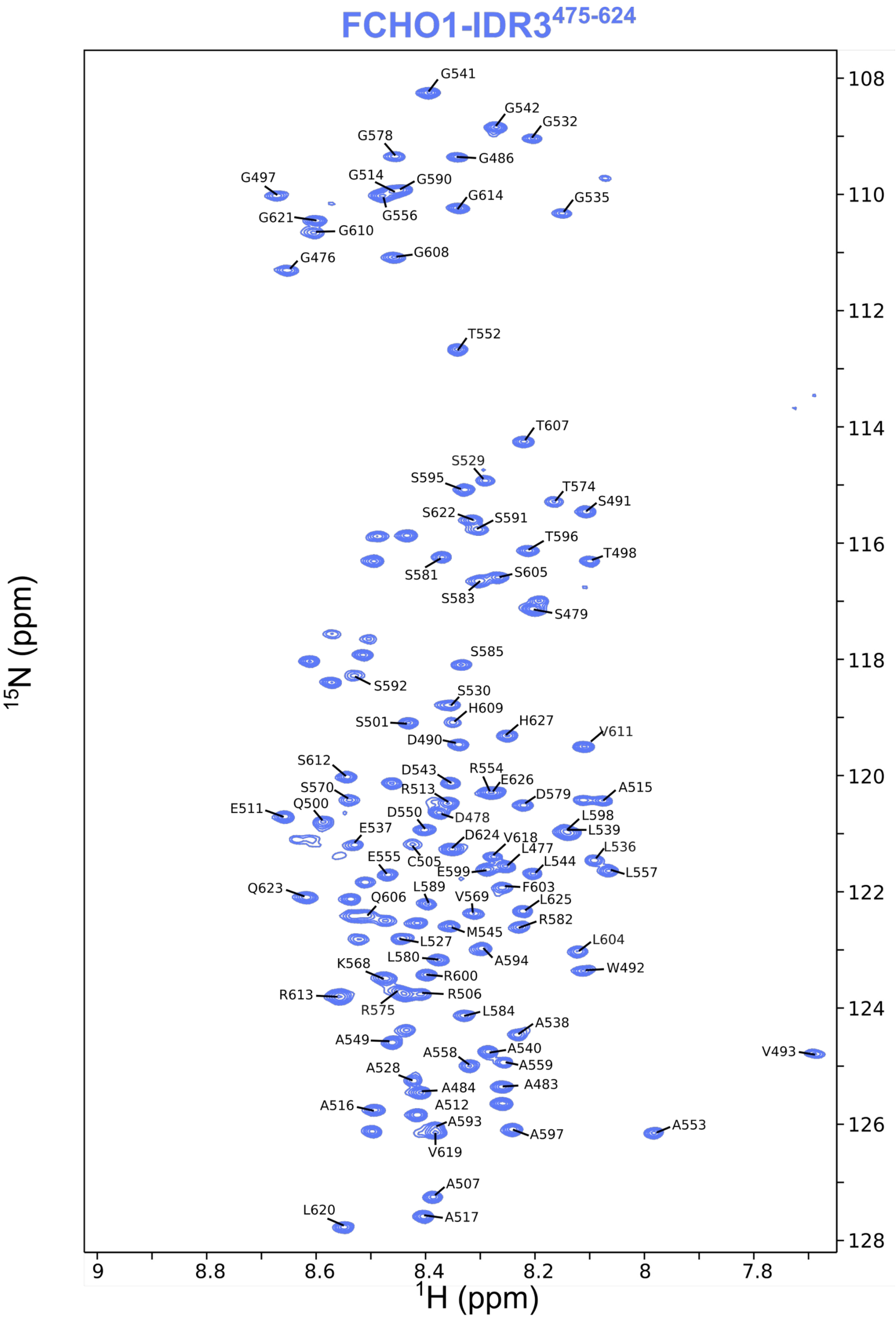


**Fig. S4. FCHO1-IDR3 assignment**. The $^{1}$H-$^{15}$N HSQC spectra of FCHO1-IDR3 construct showing the backbone resonance assignments as one letter amino acid code.

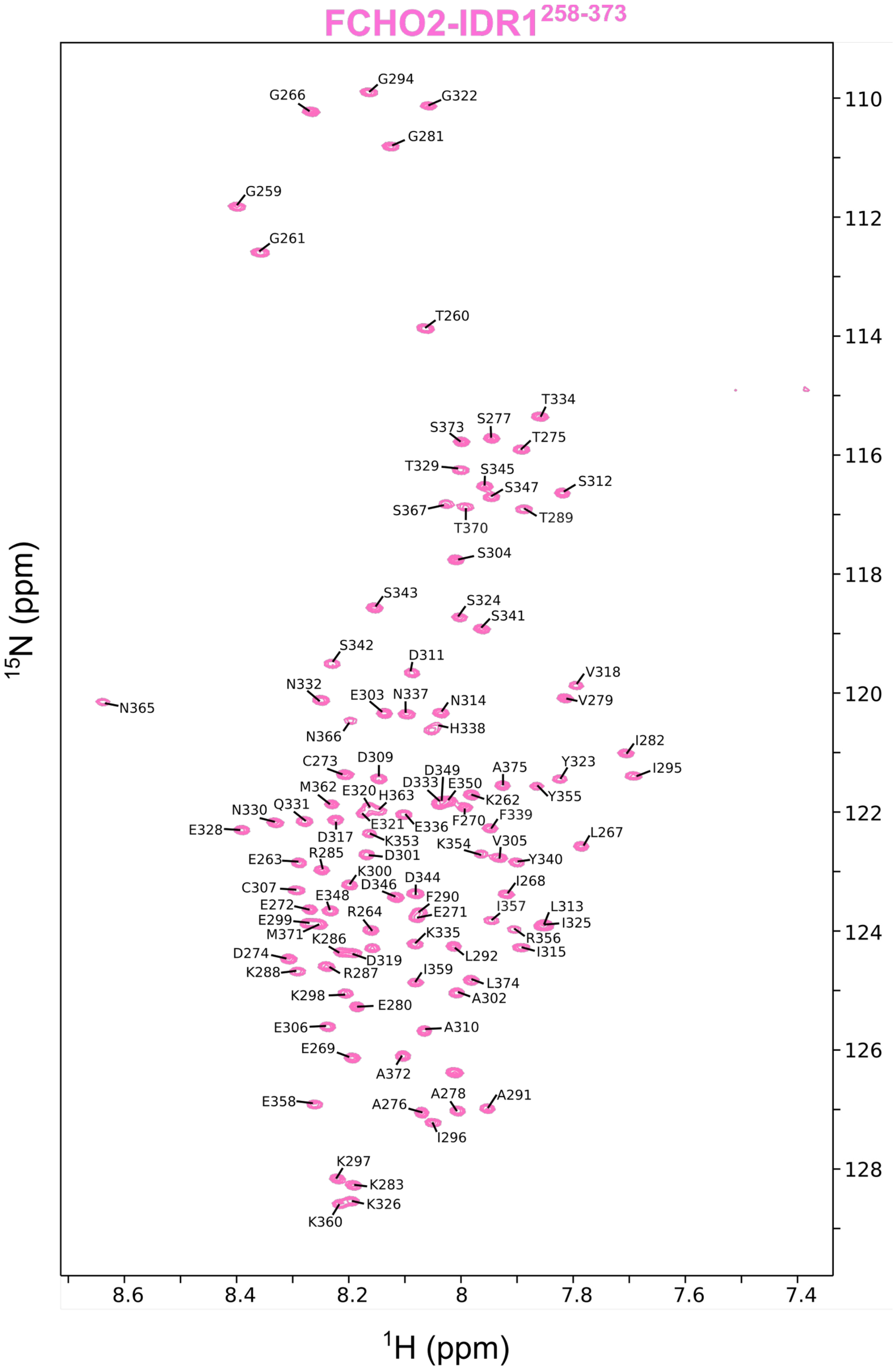


**Fig. S5. FCHO2-IDR1 assignment.** The $^{1}H$-$^{15}N$ HSQC spectra of FCHO2-IDR1 construct showing the backbone resonance assignments as one letter amino acid code.

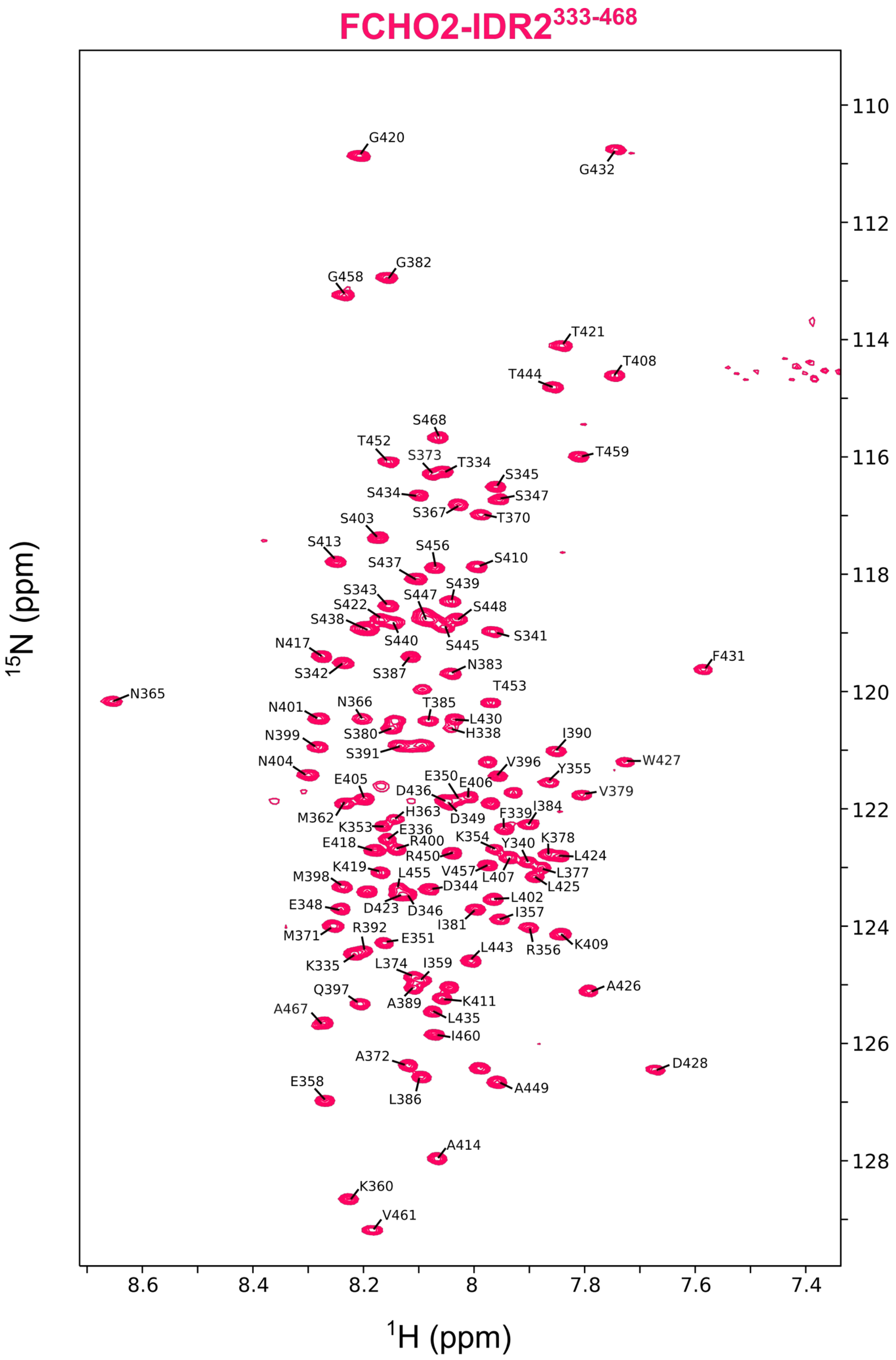


**Fig. S6. FCHO2-IDR2 assignment.** The $^{1}H$-$^{15}N$ HSQC spectra of FCHO2-IDR2 construct showing the backbone resonance assignments as one letter amino acid code.

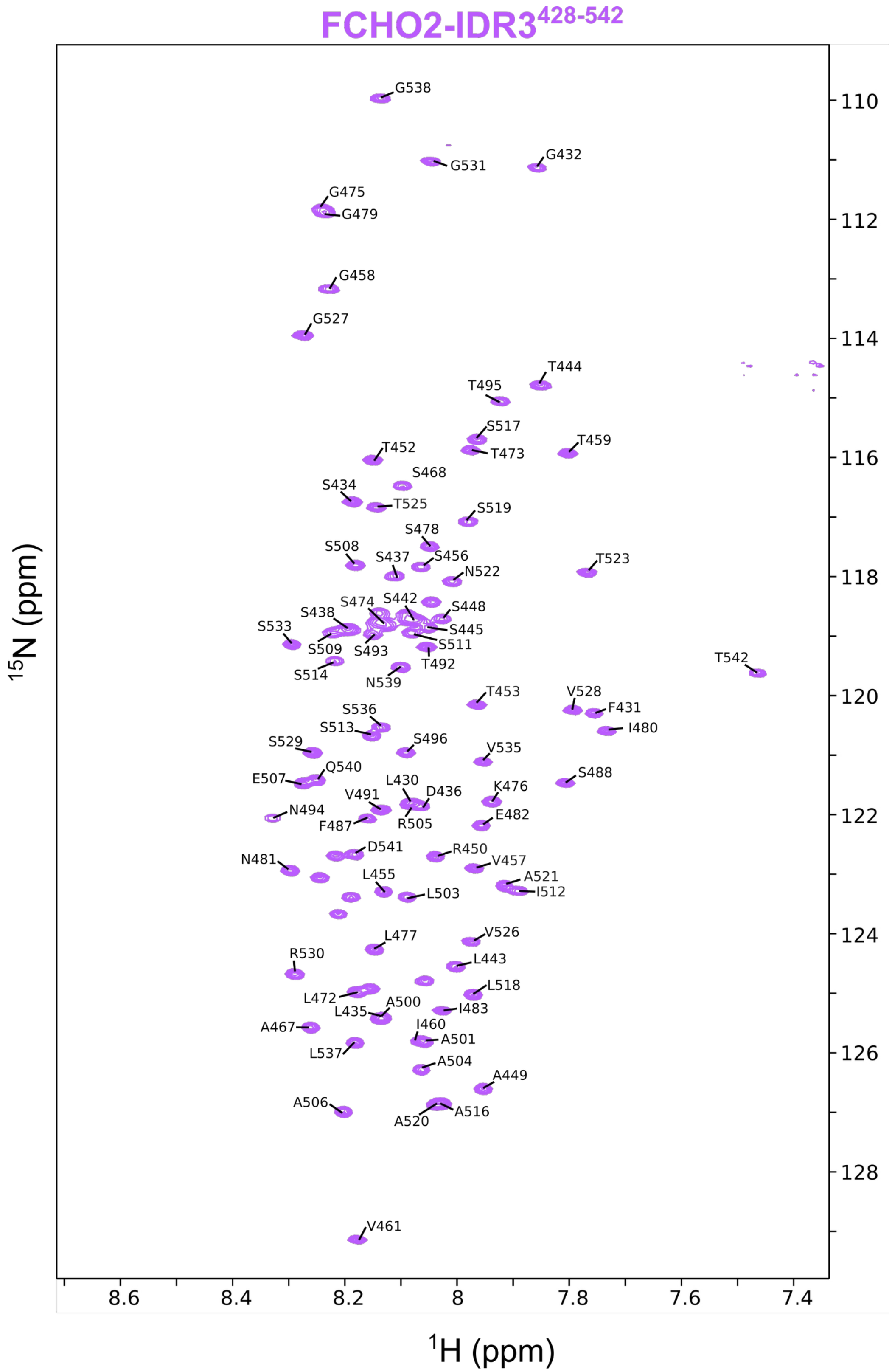


**Fig. S7. FCHO2-IDR3 assignment.** The $^1$H-$^{15}$N HSQC spectra of FCHO2-IDR3 construct showing the backbone resonance assignments as one letter amino acid code.

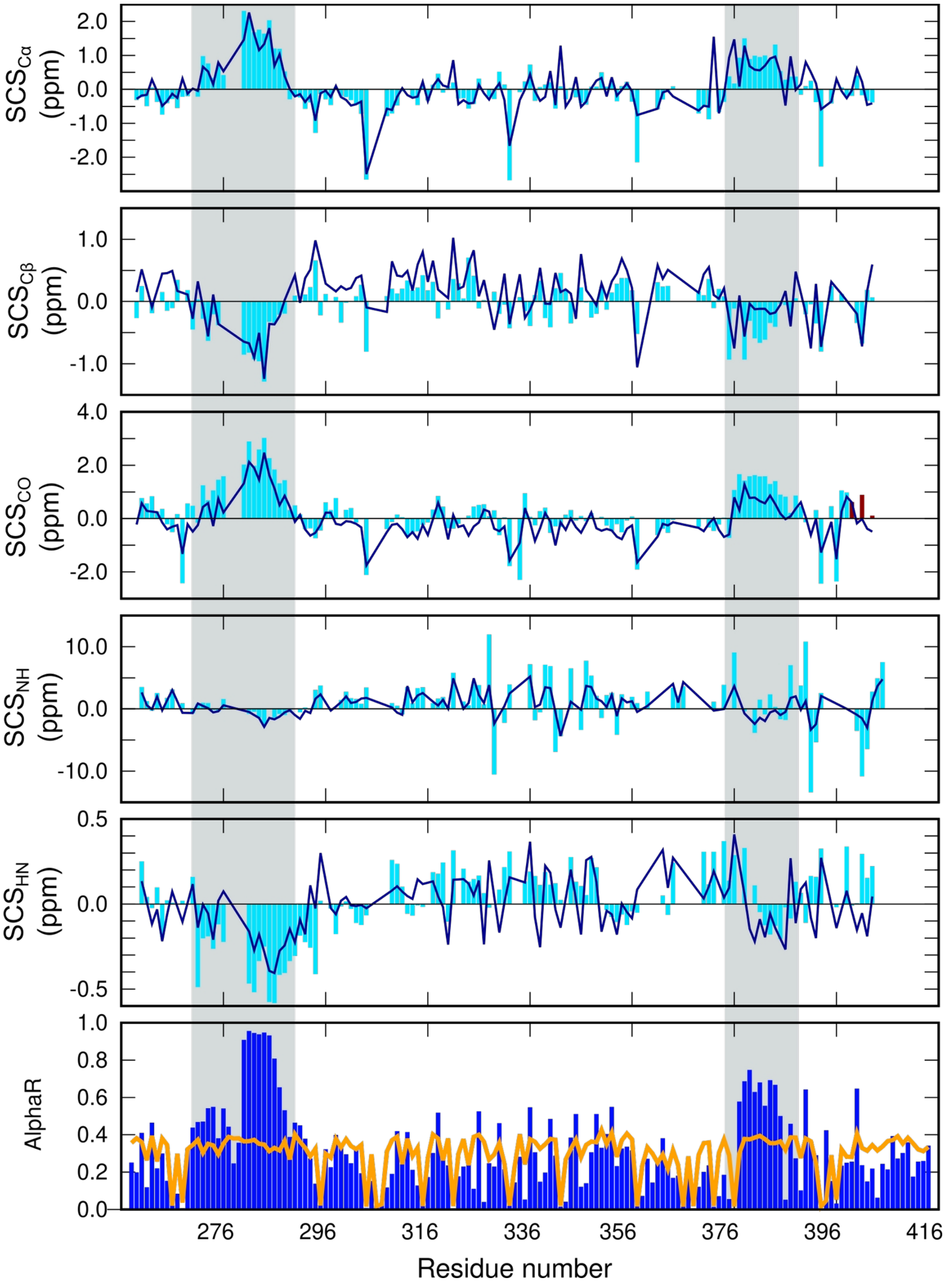


**Fig. S8. Secondary chemical shifts of FCHO1 IDR1**. SCSs (cyan) were calculated with respect to random coil chemical shifts from RefDB[43]. Back calculation from ASTEROIDS ensemble is shown as dark blue lines above the experimental values in cyan. An ASTEROIDS ensemble (bottom panel) illustrates a large increase in helical conformation (blue bars) for the two section squared in gray, as compared to random coiled (orange line).

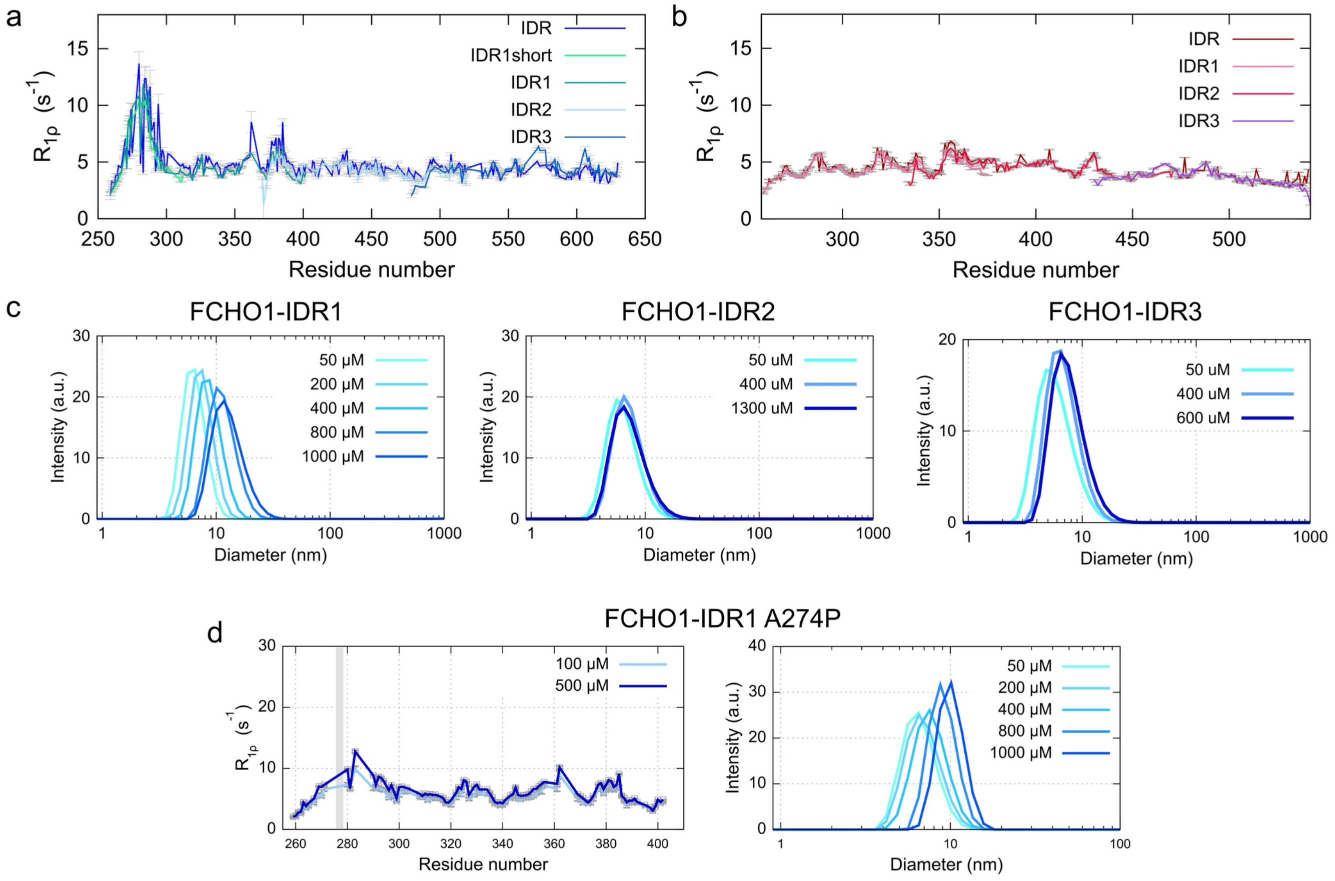


**Fig. S9. Dimerization evaluation of FCHO1 and 2 IDR short constructs. (a, b)** Overlay of $^{15}N$ $R_{1\rho}$ spin relaxation rates of FCHO1 (a) and FCHO2 (b) IDR with the short constructs used for backbone assignment (IDR1, IDR2 and IDR3). **(c)** DLS measurements of FCHO1-IDR1 short constructs upon increasing concentrations. **(d)** Self-assembly capacity of A274P FCHO1-IDR1 mutant, evaluated by $^{15}N$ $R_{1\rho}$ spin relaxation (left) and DLS (right) measurements at different concentrations. In the $R_{1\rho}$ plots, gray bar indicate the position of A→ P mutation.

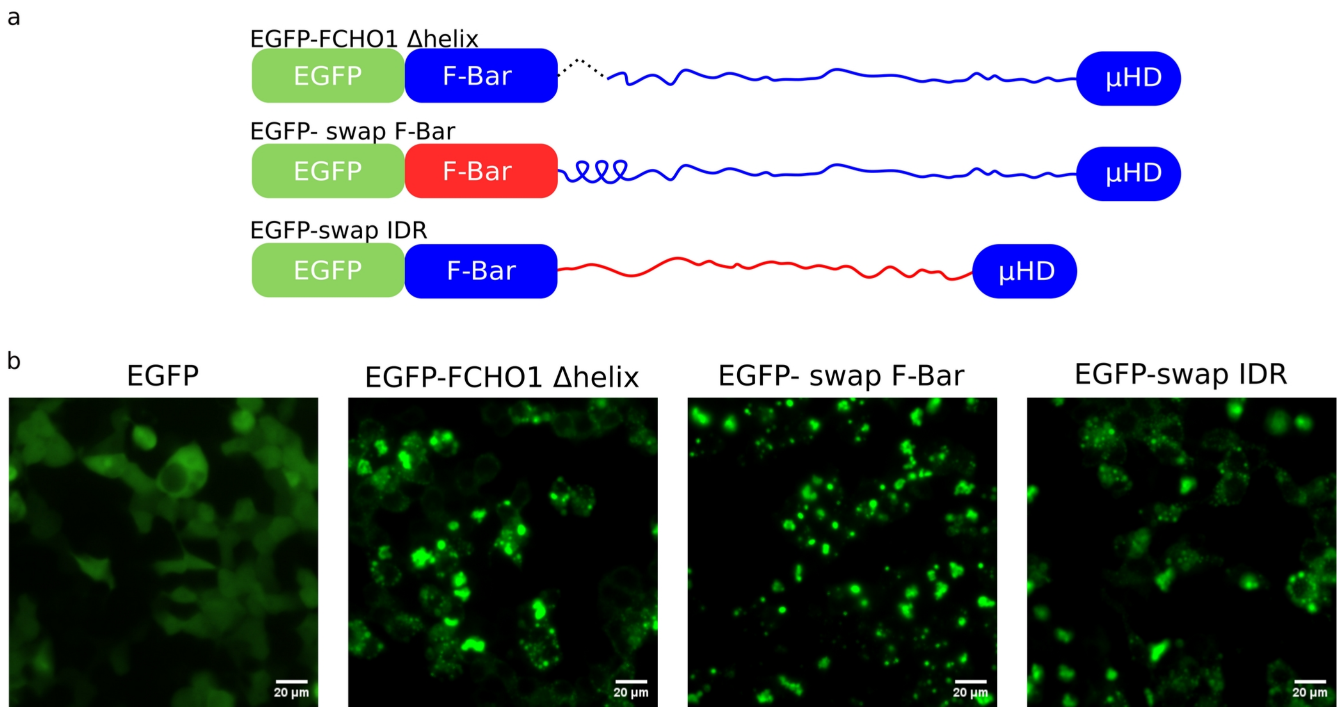


**Fig. S10. *In cell* transfection of FCHO chimeras. (a)** Cartoon representation of the full length FCHO chimera constructs fused to EGFP used in this study. **(b)** Fluorescence microscopy images of HEK cells transfected with the FCHO variants described in panel (a).

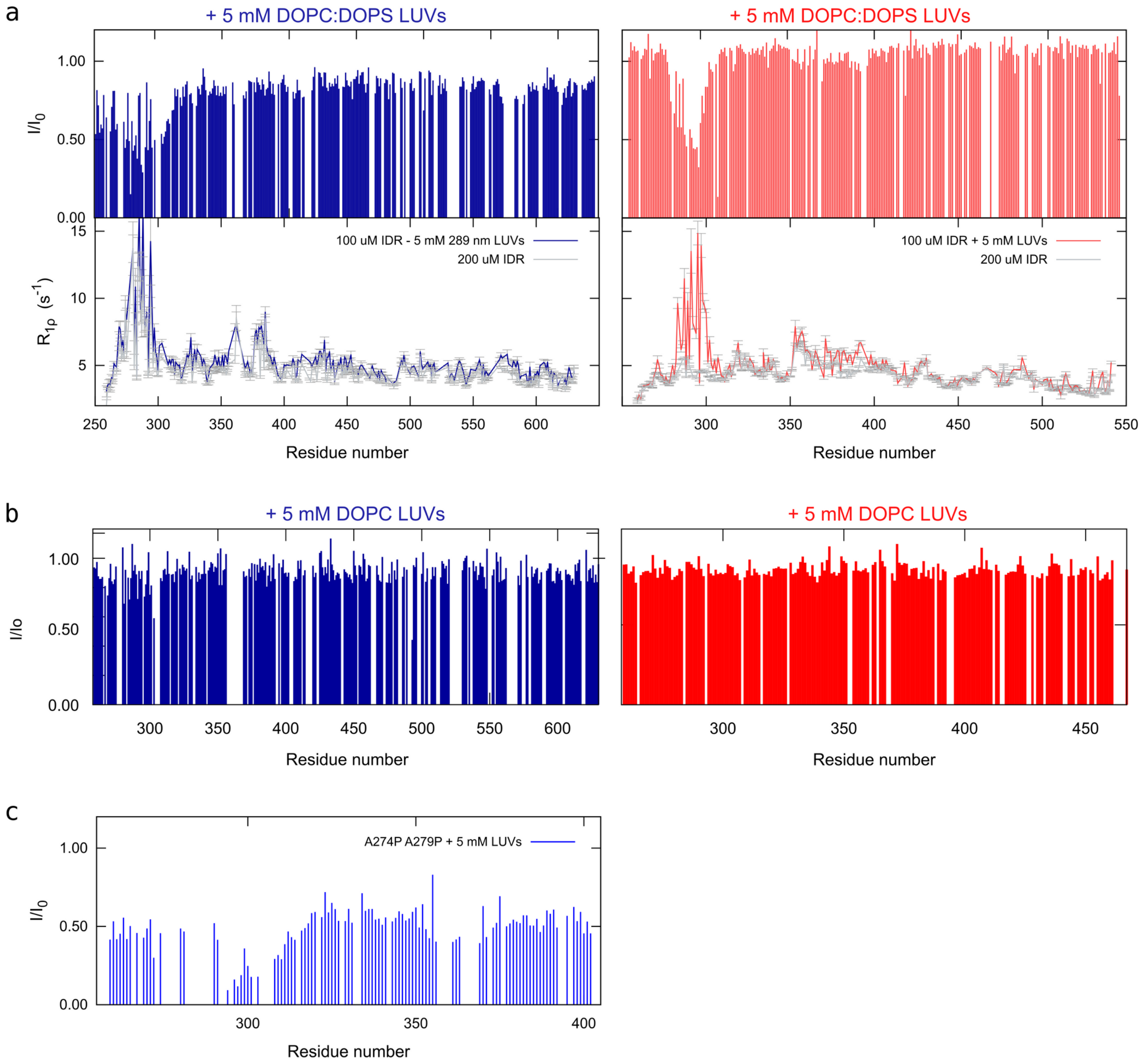


**Fig. S11. Interaction of FCHOs full IDR and mutant with LUVs**. **(a)** Intensity ratios ($I/I_0$) and $^{15}N$ $R_{1\rho}$ spin relaxation of FCHO1 (left) and FCHO2 (right) IDR in the absence and presence of 300 nm LUVs (DOPC:DOPS 1:1). **(b)** Intensity ratios ($I/I_0$) of FCHO1 (left) and FCHO2 (right) IDR in the absence and presence of 300 nm LUVs (DOPC). **(c)** Intensity ratios ($I/I_0$) of FCHO1-IDR1 A274P A279P mutant interacting with 120 nm LUVs (DOPC:DOPS 1:1).

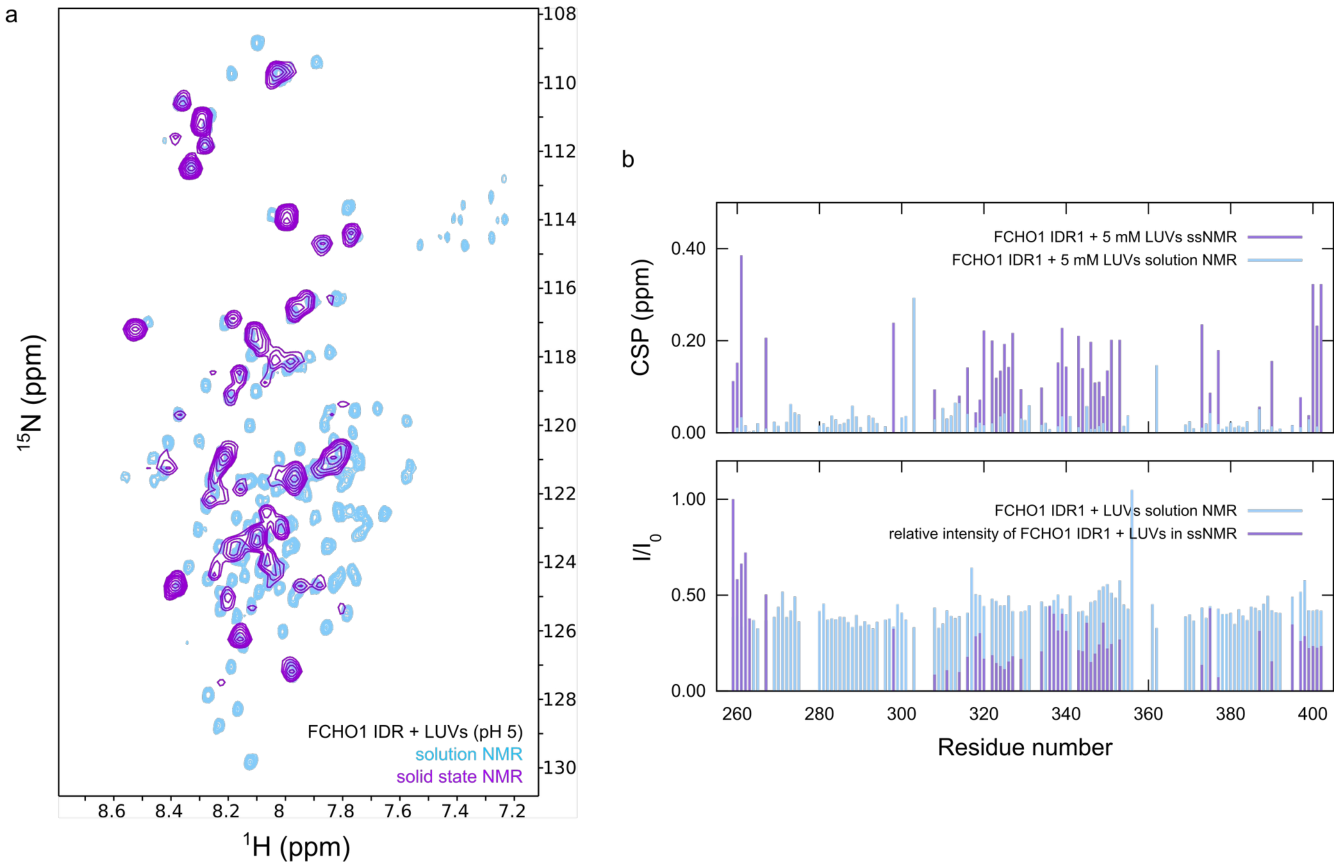


**Fig. S12. Interaction of FCHO1 IDR1 with LUVs by solid state NMR**. (a) Overlay of FCHO1-IDR1 spectra in the presence of DOPC:DOPS LUVs measured by solution (cyan) and solid state (purple) NMR. (b) CSPs and intensity ratios of FCHO1 IDR1 upon interaction with LUVs. Solid state NMR intensities are relative to the most intense signal (the N-terminal residue).

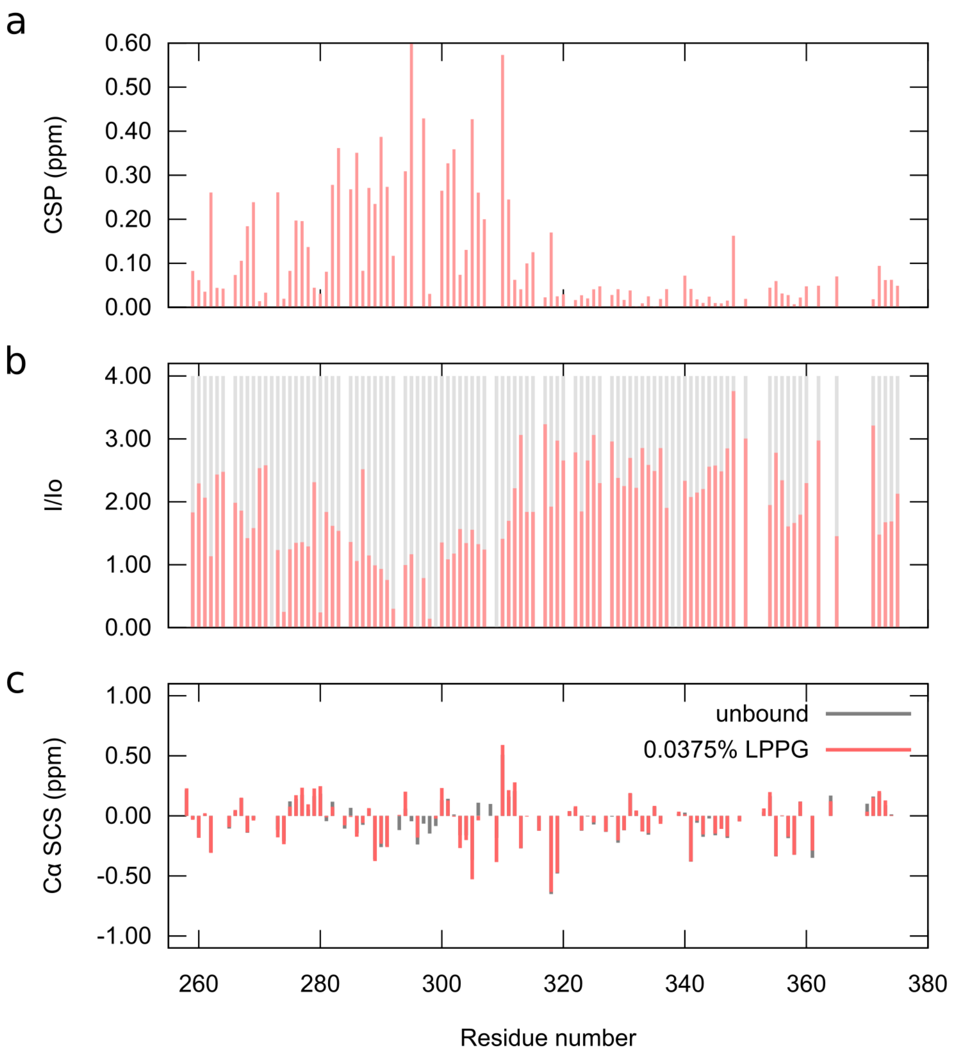


**Fig. S13. Interaction of FCHO2 IDR1 with LPPG micelles. (a,b)** Chemical shift perturbations and relative intensity ratios of FCHO2-IDR1 signals upon binding with LPPG micelles. **(c)** Cα secondary chemical shifts of FCHO2 IDR1 alone (gray) and interacting with LPPG micelles.

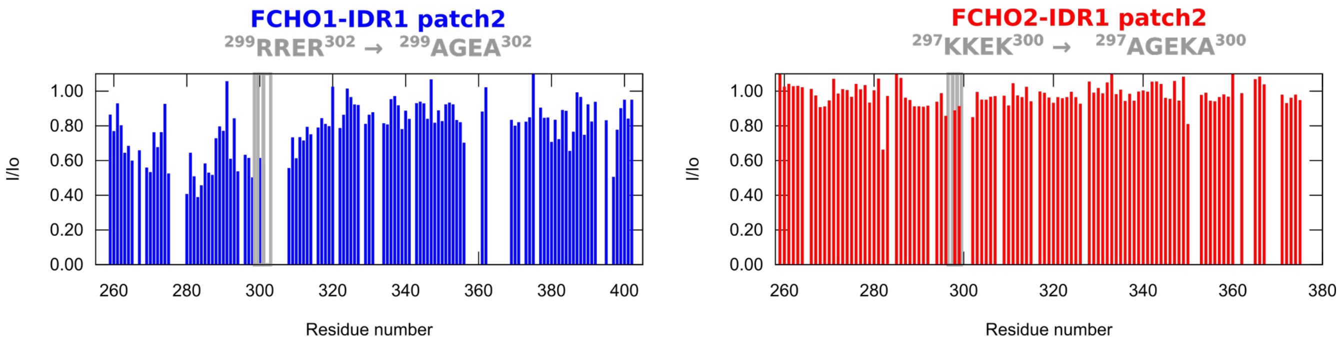


**Fig. S14. Interaction between LUVs and FCHO IDR1 patch2 mutants**. Intensity ratios ($I/I_0$) and FCHO1 (left) and FCHO2 (right) IDR1 patch2 mutants in the absence and presence of 100 nm LUVs (DOPC:DOPS 1:1).
